# Single-cell level transcriptome of the maize pathogenic fungi *cochliobolus heterostrophus* race O in infection reveal the virulence related genes, and potential circRNA effector

**DOI:** 10.1101/350785

**Authors:** Meng Wang, Shaoqing Wang, Jianan Sun, Yaqian Li, Kai Dou, Zhixiang Lu, Jie Chen

**Author notes:** These authors contributed equally to this work. Corresponding author: Jie Chen.

## Abstract

*Cochliobolus heterostrophus* is a crucial pathogenic fungus that causes southern corn leaf blight (SCLB) in maize worldwide, however, the virulence mechanism of the dominant race O remains unclear. In this report, the single-cell level of pathogen tissue at three infection stages were collected from the host interaction-situ, and were performed next-generation sequencing from the perspectives of mRNA, circular RNA(circRNA) and long noncoding RNA(lncRNA). In the mRNA section, signal transduction, kinase, oxidoreductase, and hydrolase, et al. were significantly related in both differential expression and co-expression between virulence differential race O strains. The expression pattern of the traditional virulence factors nonribosomal peptide synthetases (NPSs), polyketide synthases (PKSs) and small secreted proteins (SSPs) were multifarious. In the noncoding RNA section, a total of 2279 circRNAs and 169 lncRNAs were acquired. Noncoding RNAs exhibited differential expression at three stages. The high virulence strain DY transcribed 450 more circRNAs than low virulence strain WF. Informatics analysis revealed numbers of circRNAs which positively correlate with race O virulence, and a cross-kingdom interaction between the pathogenic circRNA and host miRNA was predicted. An important exon-intron circRNA Che-cirC2410 combines informatics characteristics above, and highly expressed in the DY strain. Che-cirC2410 initiate from the pseudogene *chhtt*, which doesn’t translate genetic code into protein. In-situ hybridization tells the sub-cellular localization of Che-cirC2410 include pathogen`s mycelium, periplasm, and the diseased host tissues. The target of Che-cirC2410 was predicted to be zma-miR399e-5P, and the interaction between noncoding RNAs was proved. More, the expression of zma-miR399e-5P exhibited a negative correlation to Che-cirC2410 in vivo. The deficiency of Che-circ2410 decreased the race O virulence. The host resistance to SCLB was weakened when zma-miR399e-5P was silenced. Thus, a novel circRNA-type effector and its resistance related miRNA target are proposed cautiously in this report. These findings enriched the pathogen-host dialogue by using noncoding RNAs as language, and revealed a new perspective for understanding the virulence of race O, which may provide valuable strategy of maize breeding for disease resistance.

**Author Summary:** The southern corn leaf blight (caused by *Cochliobolus heterostrophus*) is not optimistic in Asia, however we have limit knowledge about the infection mechanism of the dominant *C.heterostrophus* race O. We take full advantage of the ideal *C.heterostrophus* genome database, laser capture microdissection and single-cell level RNA sequencing. Hence, we could avert the artificial influence such as medium, and profile the real gene mobilization strategy in the infection. The results of coding RNA section were accessible, virulence related genes (such as the signal transduction, PKS, SSP) were detected in RNA-seq,which accord with previous reports. However, the results of noncoding RNA was astonished, 2279 circular RNAs (circRNA) and 169 long noncoding RNAs (lncRNA) were revealed in our results. Generally, the function of noncoding RNA was hypothesized in single species, but we boldly guess that the function of circRNA is rather complicated in the pathogen-host interaction. Finally, the circRNA in-situ hybridization (ISH) demonstrate the secretion of pathogen circRNA into the host tissue. By bioinformatic prediction, we found a sole microRNA target, and proved the interaction between circRNA and microRNA. These findings are likely to reveal a novel pathogen effector type: secreted circRNA.

## Introduction

Southern corn leaf blight (SCLB), which is caused by *Cochliobolus heterostrophus* (*C. heterostrophus*), is a major fungal disease on maize worldwide(1, 2) (3) (4). T-toxin is the lethal effector which target on URF13 of T-cytoplasmic male-sterile(TCMS) maize line (5),(6),(7). While, the effectors of the dominant race O and the targets of host maize remain unclear(8). Fortunately, the released race O genome database(8),(9), laser capture microdissection(LCM)(10–12) and isothermal-linear amplification(13),(14),(15) enable us to profile the whole transcriptome (including coding mRNAs and noncoding circRNAs, lncRNAs) from a few single-cell level mycelium debris. Methology above are helpful for revealing the true genome mobilization strategy in infection progress, and averting the man-made interference such as artificial medium.

mRNA-sequencing is the frequently used way after LCM(16, 17), while increasing reports have disclosed that noncodingRNAs play numerous roles in eucaryon (18–20), and regulate pathogen-host interaction(21–23). For example, miR393 is a conserved microRNA triggered by PAMPs, which represses F-box auxin receptors and consequently impedes auxin signaling(24). *B. cinerea* small RNAs bind to the AGO1 of *Arabidopsis thaliana* and *Solanum lycopersicum*, and then selectively silence the host immunity genes(25). However, the role of circRNA or lncRNA in pathogen-host interaction remains unknown.

This study aimed to understand the *C.heterostrophus* race O infection on the basis of mobilizing the full potential of genome. Not only mRNA but also circRNA and lncRNA were successfully profiled from infecting pathogen of single-cell magnitude. It is found that race O transcribe amount of noncoding RNAs(especially circRNA) in infection, and a cross-kingdom interaction between pathogenic circRNA and host microRNA is revealed.

## Results

### 1. Sampling and transcriptome data size of *C.heterostrophus* race O

Race O high virulence strain DY and low virulence strain WF were tagged with green fluorescent protein (GFP). Both GFP-tagged strains retained their idifferential nfection capability on the maize inbred lines B73 (susceptible to SCLB) and Mo17 (resistant to SCLB) (Fig. 1C). Three distinguishable stages of the *C. heterostrophus* race O during the infection process were introduced for analysis: (1) conidium (CO, 0-0.5 hpi, Fig.1D, Fig.1F), (2) scattered mycelium (SM, 9-11 hpi, Fig.1E, Fig.1G), and (3) dense bundles of thick mycelium (DM, the mycelium exhibited tropism to motor cells, 60-72hpi, Fig.1H, Fig.1I, S1 Fig.). In each sample, 10 ng total RNA collected from infection situ was amplified to cDNA library (S2 Fig.), at least 7 GB (200 folds depth comparing to the genome size) filtered data were obtained, which reserve the mRNA, circRNA and lncRNA information (S1_table).

**Figure 1.**
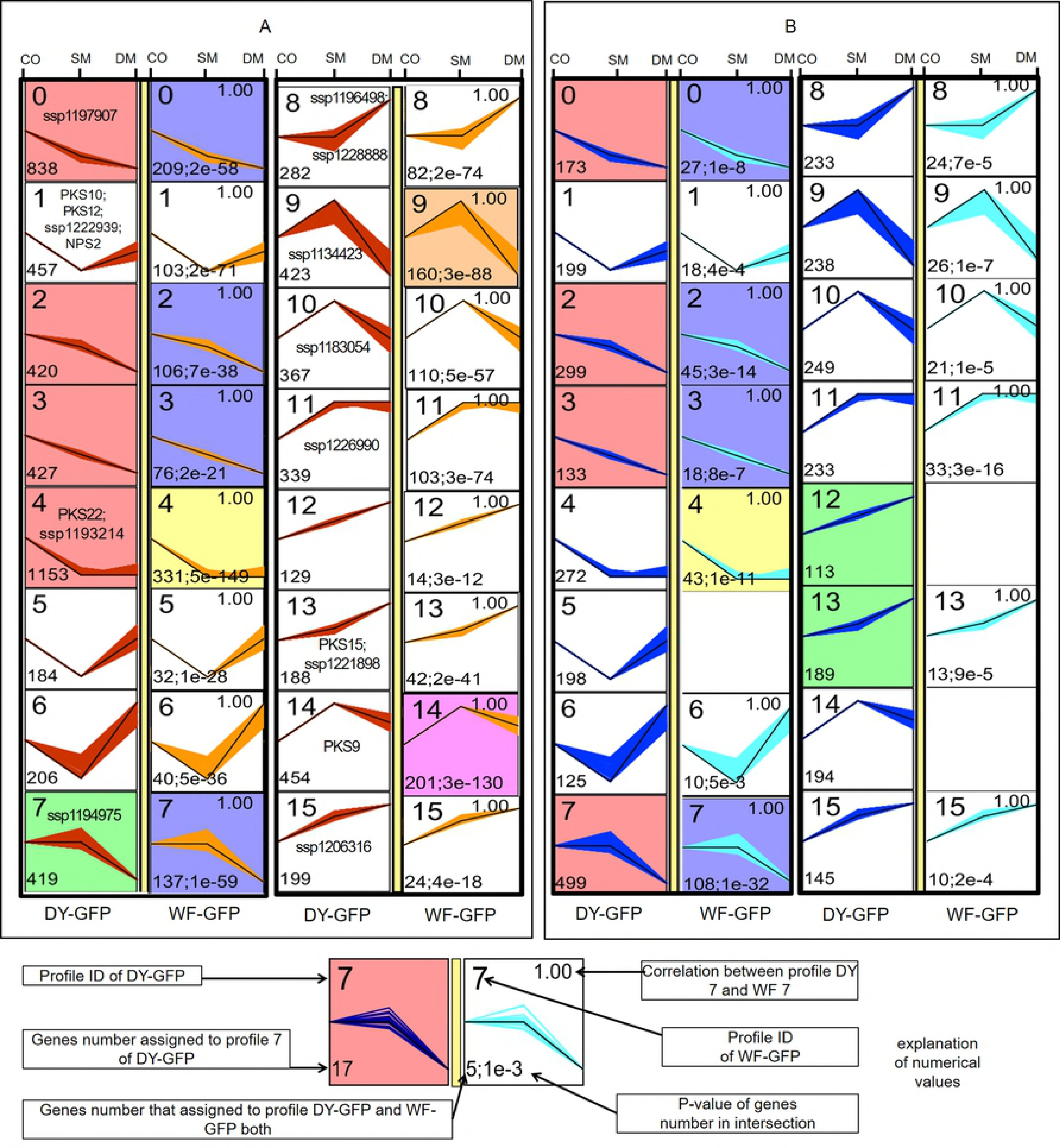
Colony, infectin stage of *Cochliobolus heterostrophus* race O strains. (A) Colony of the high-virulence strain DY-GFP on plate. (B) Colony of the low-virulence strain WF-GFP on plate. (C) Infection of DY-GFP and WF-GFP on two the *Zea mays* inbred lines B73 and Mo17. The left side of the foliage vein was inoculated with WF-GFP conidia at the concentration of 1×10^5^ spores/ml. The right side of the vein was inoculated with DY-GFP conidia at the same concentration. B73 exhibited susceptibility to the pathogen, whereas Mo17 exhibited resistance. (D) Conidium stage of DY-GFP; the timing was 0 hpi. (E) Scattered mycelium stage of DY-GFP; the timing was 9 hpi. (F) Conidium stage of WF-GFP; the timing was 0 hpi. (G) Scattered mycelium stage of WF-GFP; the timing was 9 hpi. (H) Dense bundles of thick mycelium stage of DY-GFP; the timing was 72 hpi. (I) Dense bundles of thick mycelium stage of WF-GFP; the timing was 72 hpi. (J) The surface structure of the B73 leaf infected by DY-GFP. (K) The surface structure of the B73 leaf infected by WF-GFP. H and I were photographed under an RFP/GFP fluorescence dual channel, whereas J and K were photographed under a bright-field channel. ST, stoma; EC, epidermal cell; MC, motor cell; DM, dense bundles of thick mycelium. Scale bar =100 μm.

### 2. Co-expression of mRNA in race O high/low virulence strains

For understanding the genome mobilization strategy of each infection stage, the co-expression mRNA in both strains were analyzed using series test of cluster (STC) (26). All mRNA transcripts were sorted into 15 profiles (Fig. 2A), and exhibited 4 kinds of expression tendency: STC profile 0, 1, 2, 3, 4 decline continuously (DC) from CO till DM stage; STC profiles 5, 6 and 8 exhibited down-regulation at the SM and up-regulation at the DM stage (DSUD); STC profiles 7, 9, 10 exhibited up-regulation at the SM and down-regulation at the DM stage (USDD); STC profiles 11-15 increase continuously (IC) from CO till DM stage.

**Figure 2.**
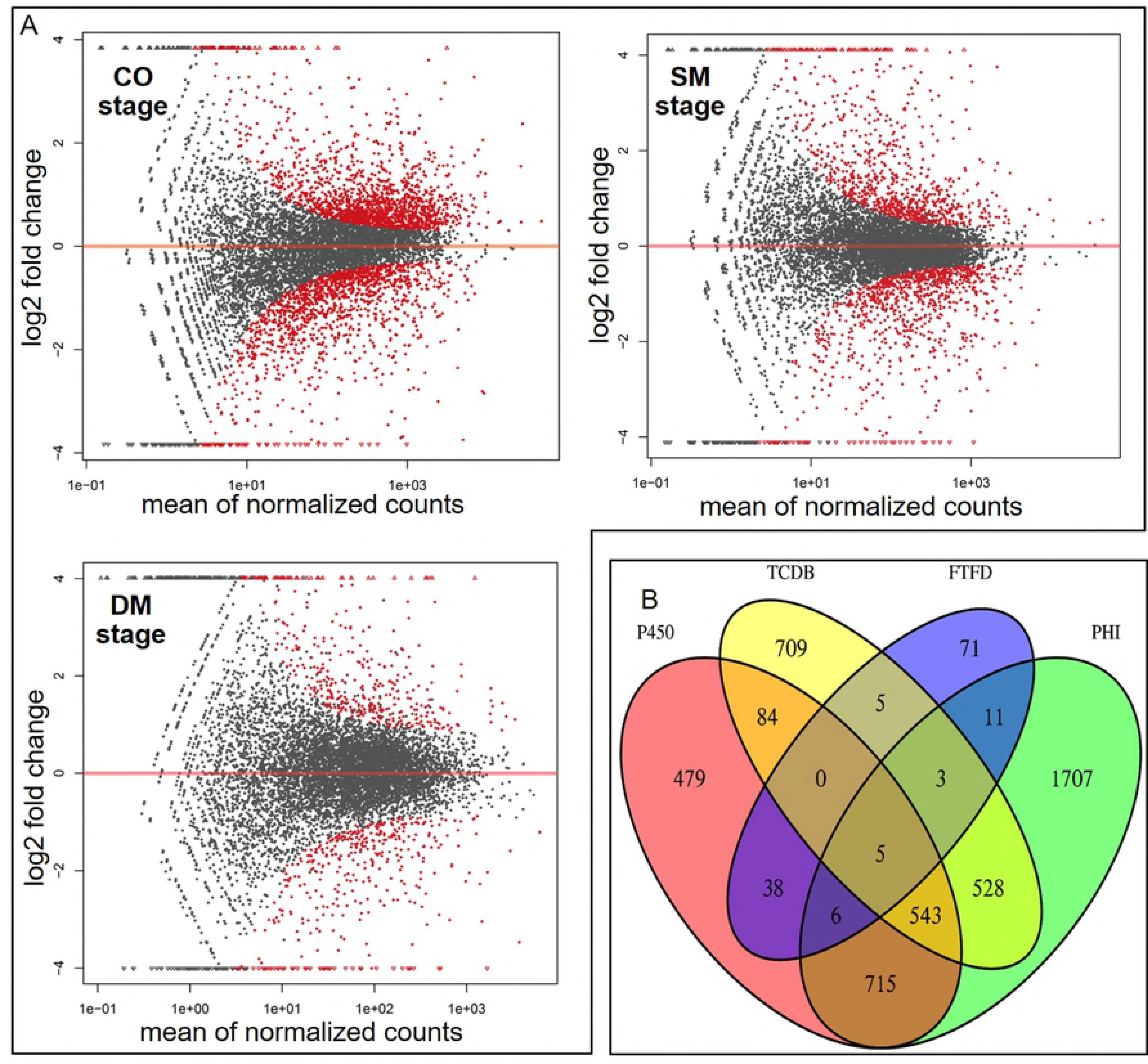
Series test of cluster (STC) analysis to present the genes that share the same expression laws in the three stages. (A) STC analysis of the mRNA transcripts. (B) STC analysis of the circRNA transcripts. The DY profiles are presented at the left side of the yellow bar, and the WF profiles are presented at the right side of the yellow bar. Annotations for numerical values are indicated by arrows at the bottom of the figure.

Significant genes of each profile were annotated with molecular function (MF) and biological progress (BP). The MF of DC profiles were enriched of nucleotide binding, histidine kinase, serine/threonine kinase, two component sensor, signal transducer, et al.; the significant BP of DC were peptidyl-histidine phosphorylation, protein modification, two-component signal (phosphorelay), phosphorylation, DNA repair, signal transduction, transport, regulation of transcription. The DSUD profiles` MF were enriched of oxidoreductase, hydrolase, dehydrogenase, catalytic, et al.; the significant BP of DSUD were carbohydrate metabolic process. The USDD profiles` MF were enriched of oxidoreductase, dehydrogenase, catalytic, protein dimerization, transporter, FAD binding, pyridoxal phosphate binding,et al.; the significant BP of USDD were ATP synthesis coupled electron transport, protein transport, transport, DNA repair, metabolic process, transport, nitrogen compound metabolic. The IC profiles’ MF were enriched of hydrolase, proteasome endopeptidase, catalytic, N-acetyltransferase, structural constituent of ribosome,et al.; the significant BP of IC were tRNA aminoacylation for protein translation, translation, carbohydrate metabolic process, metabolic process, protein modification.

### 3. Differential-expression of mRNA in race O high/low virulence strains

Co-expression analysis is helpful for disclosing the major pathogenicity related function of each infection stage, besides this, we were curious about the genes that related to pathogenicity differentiation. Generally, the up-regulated mRNAs of DY were not as abundant as WF (even though the virulenc of DY is high), especially at the CO stage (Table 1). The number of differential mRNAs decreased, however the average expression level expand along with infection progress (Fig. 3).

**Table 1.** Differential expression RNAs of race O high/low virulence strains

| infection stage | mRNA up-regulated |  | lncRNA up-regulated |  | circRNA up-regulated |  |
| --- | --- | --- | --- | --- | --- | --- |
|  | DY | WF | DY | WF | DY | WF |
| CO | 548 | 763 | 17 | 31 | 469 | 431 |
| SM | 527 | 512 | 21 | 26 | 513 | 144 |
| DM | 399 | 393 | 17 | 11 | 175 | 132 |

**Figure 3.**
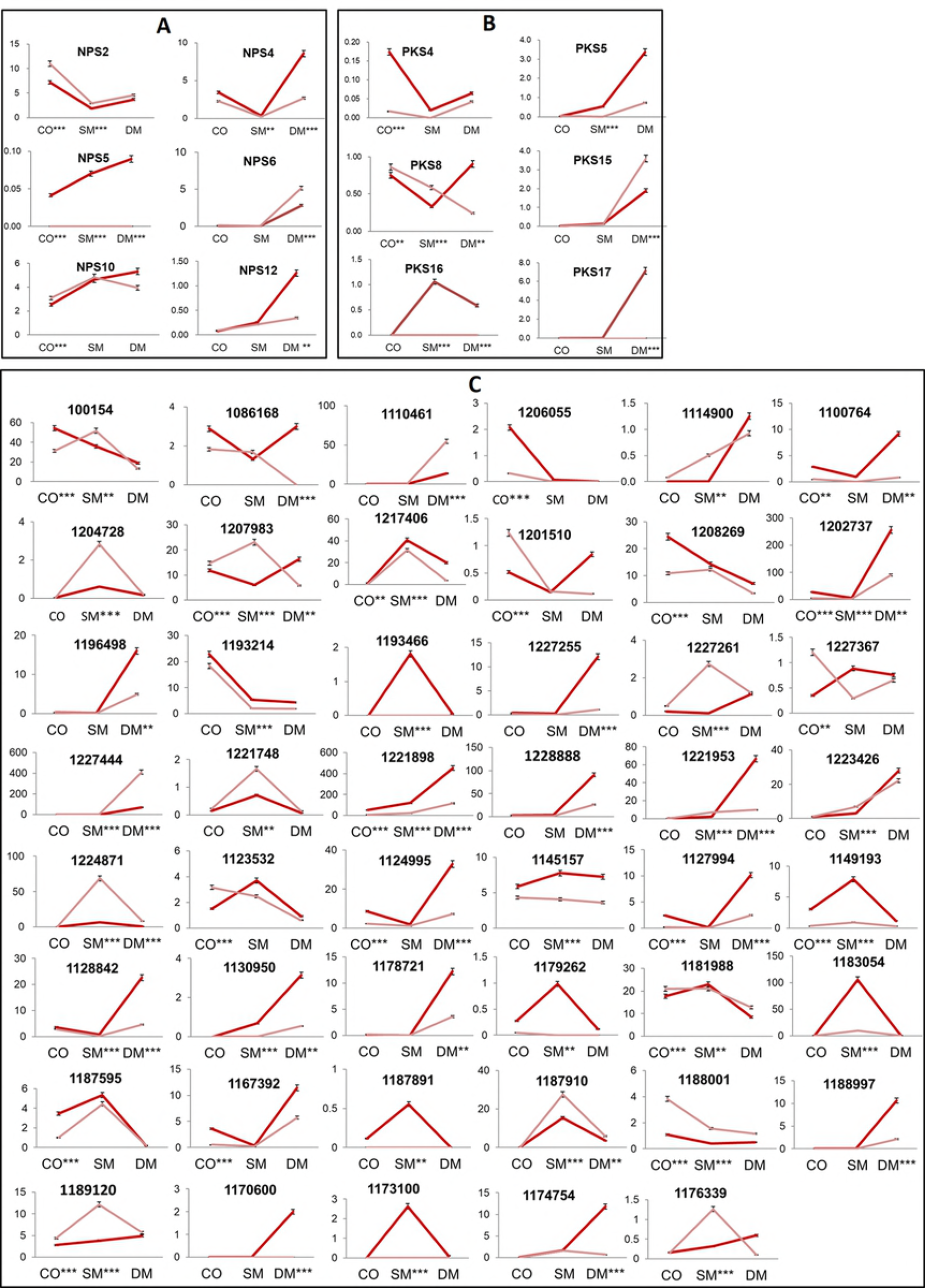
The mRNA differential status between high-virulence DY and low-virulence WF. (A) MA diagram for displaying the differential genes of each infection stage, the Y-axis represent the log2 fold change, the X-axis represent the gene mean normalized counts, the red spots mean that the differential is significant (P<0.05). (B) the venn diagram for displaying genes that fitted in the P450, TCDB, FTFD and PHIdatabase.

Annotation with molecular function(MF) was used for analysing the differential mRNAs` functions. At the CO stage, DY significantly up-regulated signal transduction compounds mainly (e.g., phosphofructokinase, GTPase activator, protein serine/threonine phosphatase, phosphatidylinositol phosphate kinase, and protein-tyrosine kinase); besides signal transduction, diverse functional genes up-regulated in WF CO stage (eg., D-arabinitol 2-dehydrogenase, transcription factor, amino acid transmembrane transporter, F420-2 glutamyl ligase, ribosomal S6-glutamic acid ligase, two-component sensor, and protein histidine kinase). At the SM stage, DY significantly up-regulated oxygenases (e.g.,monooxygenase, dioxygenase, and fluorene oxygenase), carboxypeptidases, hydroxylase, and dehydrogenase; at the WF SM stage, the up-regulate redox genes (eg., ferric-chelate reductase) were not as abundant as DY, we also found DNA helicase, RNA helicase, carbon-sulfur lyase, laccase, endo-1,6-alpha-mannosidase, et al. ․ At the DM stage, DY significantly up-regulated serine carboxypeptidase, procollagen-proline dioxygenase, procollagen-proline 4-dioxygenase, carboxypeptidase D, vacuolar carboxypeptidase, carboxypeptidase C, polygalacturonase, serine-type peptidase, amino acid transmembrane transporter, subtilase, cutinase, oxidoreductase, L-aminoadipate-semialdehyde dehydrogenase, hydrolase, F420-2 glutamyl ligase, chitin binding; WF also up-regulated oxidoreductase, monooxygenase, hydrolase. The other up-regulated genes in WF DM stage were inorganic phosphate transmembrane transporter, L-arabinose isomerase, beta-glucosidase, sugar:hydrogen symporter, NADH dehydrogenase, tetracycline:hydrogen antiporter, choline dehydrogenase, carbon-nitrogen ligase.

On the whole, both co-expression and differential-expression analysis revealed that the signal related genes were mostly significant at the CO stage, redox related genes were mostly significant at the SM stage, metabolic related genes were mostly significant at the DM stage.

### 4. Expression laws of the classical virulence factors

NPSs, PKSs and SSPs are three important gene catalogs related to the virulence of *C.heterostrophus*(9), and their expression laws of infection were monitored. NPS2(CocheC5_1.BGT_BGT_e_gw1.33.2.1) was found in STC1 (Fig. 2), which highly expressed at the CO and DM stage, down-regulated at the SM stage. Meanwhile, the NPS2 expression level of DY at the CO stage were 0.51 fold (P=0.000) of WF, and 0.64 fold (P=0.003) of WF at the SM stage(Fig. 4A). The expression of NPS4 (e_gw1.3.1152.1), NPS5 (e_gw1.5.371.1) and NPS12 (CocheC5_1.estExt _fgenesh1_pm.C_50043) were higher in DY strain compared to WF (Fig. 4A), the expression of NPS6 (CocheC5_1.BGT_estExt_fgenesh1_pg. C_250126) and NPS10 (estExt_Genewise1Plus.C_8_t10324) were lower in DY strain (Fig. 4A). PKS22 (in STC4) exhibited decline continuously tendency in both strains, PKS 15 (in STC13) and PKS9 (in STC14) exhibited increase continuously tendency of both strains (Fig. 2A). Although PKS15 co-expressed in both strains, DY was with lower expression level at the DM stage. There were 5 PKS, which highly expressed in DY strain (Fig. 4B). SSPs is the most abundant catalog: a number of 10 SSPs were found with co-expression laws (Fig. 2A), and 47 SSPs differentially expressed (Fig. 4C). The deploy of NPSs, PKSs and SSPs in infection stages were related to the universal virulence factor and pathogenicity differentiation of race O.

**Figure 4.**
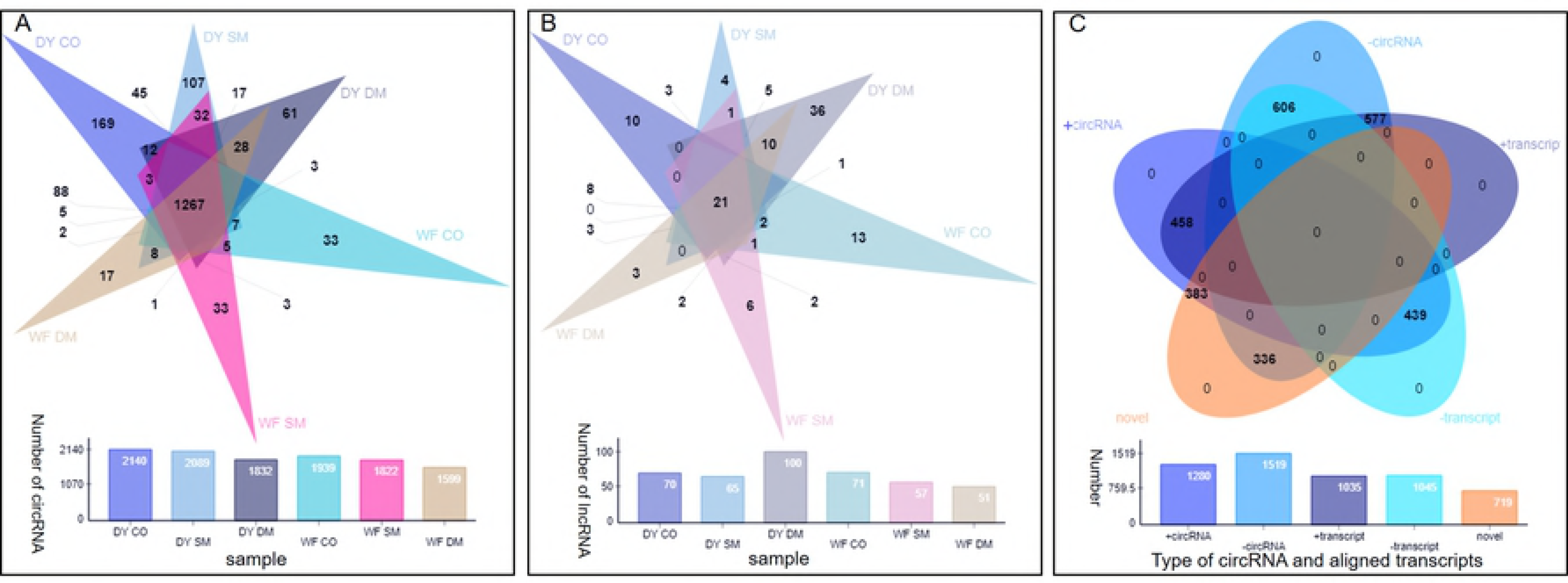
The expression of classical virulence related genes. (A) The differential NPS between *C.heterostrophus* race O strains DY and WF (differential significance P<0.05). (B)The differential PKS between race O strains DY and WF (differential significance P<0.05). (C) The differential SSP between race O strains DY and WF (differential significance P<0.05).

We also blast the co-expression and differential-expression genes in the fungal transcription factor database (FTFD)(27), transporter classification database (TCDB)(28, 29), pathogen-host interactions database (PHI)(30) and cytochrome P450 (P450)(31). In the co-expression analysis, 17 matched genes were fitted in PHI database (S2_table), whose functions were related to monooxygenase, ankyrin, ferric reductase, GMC oxidoreductase, cullin-RING ubiquitin ligase, NPS2, ABC transporter, et al. ․ In the differential expression analysis, 1870 genes were fitted in the P450 base, 1877 genes were fitted in the TCDB base, 139 genes were fitted in the FTFD base, 3518 genes were fitted in the PHI base. There are numberous genes which intersect in multiple database, which tell the complication of virulence related gene function (Fig. 3B).

### 3. Genome-wide noncoding RNA transcription during *C.heterostrophus* race O infection

A total number of 2799 circRNAs and 169 lncRNAs were filtered from sequencing data (FPKM>1) (S3_table), and all the lncRNAs are novel transcripts. Pathogens transcribe more noncoding RNAs at the earlier infection stage, except lncRNA of DY DM stage (Fig. 5). There are 1267 circRNAs and 21 lncRNAs intersected in all samples(Fig. 5). 97.89% circRNAs` length are shorter than 2000bp, 744 circRNAs` length are between 122-222bp, 634 circRNAs’ length are between 222-322bp, and 137 circRNAs` length were longer than 3000bp (Fig. 6B). The shortest circRNA is 122bp ( circRNA_2652, circRNA_5425), and the longest circRNA is 47910bp ( circRNA_1028) (S3_table). Multiple circRNAs were aligned to same one transcript, for example, 124 circRNAs were aligned to the transcript of gene e_gw1.9.987.1(function unknown), and the length distributed from 128bp to 1852bp.230 circRNAs were aligned to the gene estExt_fgenesh1_pm.C_90390 (function unknown), and the length distributed from 126 bp to 4752bp (S3_table).

**Figure 5.**
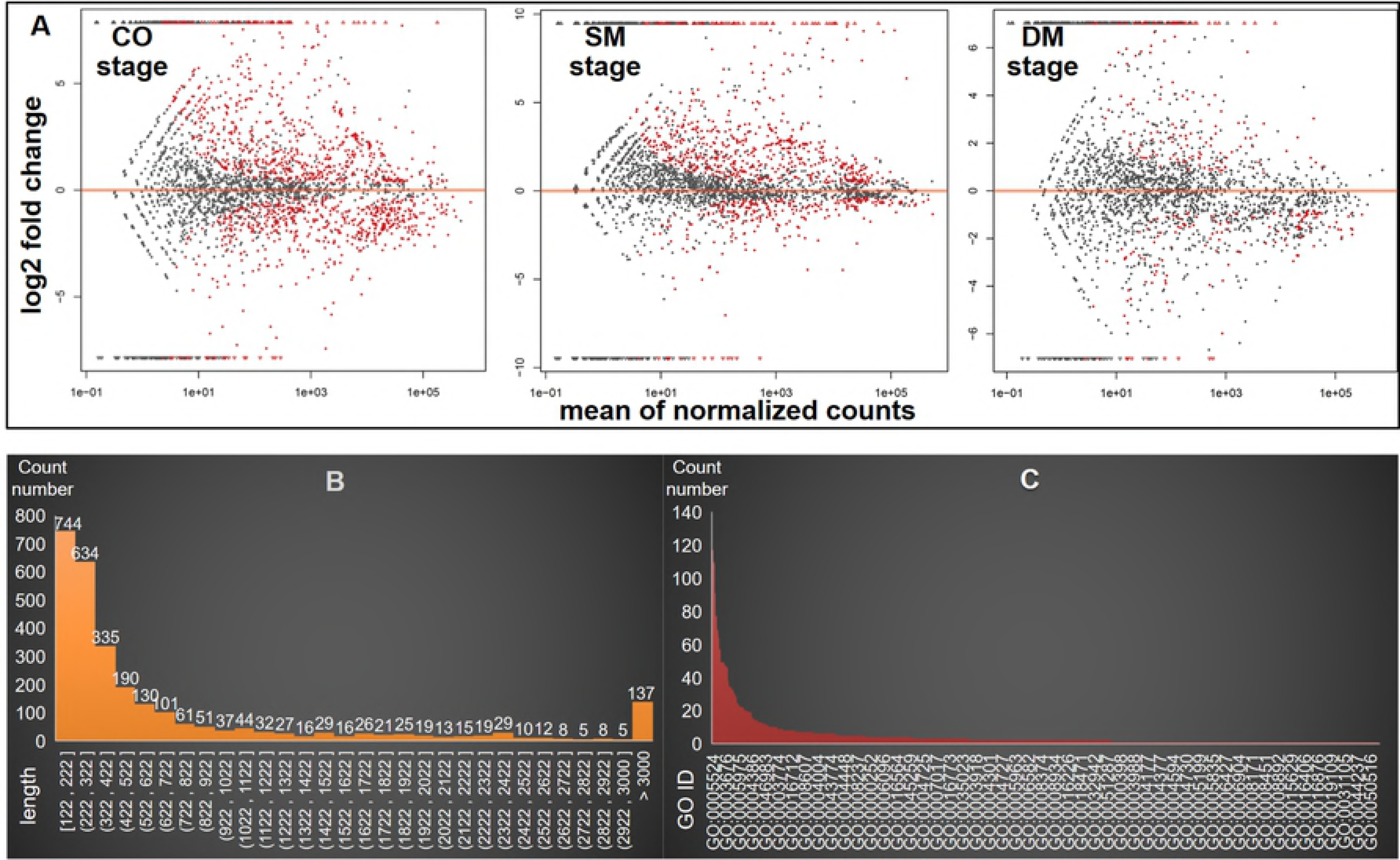
The statistical information of noncoding RNAs. (A) The number of circRNA in each infection stage. (B) The number of lncRNA in each infection stage. (C) The strand statistics of circRNA. The intersected number was demonstrated by venn diagram, and the number of each sample was demonstrated by column.

Sense-antisense RNA were widely revealed in the transcriptome analysis, and display regulation function in eukaryote(32–34).The *C.heterostrophus* race O circRNAs and the aligned linear transcripts were also found asymmetric-transcription: 1280 circRNAs were intended to be sense-strands, and the linear transcripts aligned in genome database divided into 458 sense-strands, 439 antisense-strands, and 383 noval transcripts. The other 1519 antisense-strand circRNAs were aligned to 577 sense-strand transcripts, 606 antisense-strand transcripts and 336 noval transcripts (Fig. 5C). Sense and antisense RNA are generally involved in RNA interference(35–37), sense-antisense RNA was also found in the *Aspergillus oryzae*(38) and fission yeast(39), and the role of pathogenic fungisense-antisense circRNA remains mystery.

A total number of 396 circRNAs shared co-expression laws between strain DY and WF, which were sorted into 15 profiles by the STC analysis (Fig. 2B). There were also 4 kinds of expression tendency in both strains: STC profile 0, 1, 2, 3, 4 declined continuously (DC) from CO till DM stage; STC profiles 6 and 8 down-regulated at the SM and up-regulated at the DM stage (DSUD); STC profiles 7, 9, 10 up-regulated at the SM and down-regulated at the DM stage (USDD); STC profiles 11, 13,and 15 increased continuously (IC) from CO till DM stage. However, strain WF lacked two profiles corresponding to profile 5 and 12 of strain DY.

Between two strains, DY transcribed around 200 more circRNAs than WF at each stage. 900, 657 and 307 differential circRNAs were revealed at the CO, SM and DM stage separately (P < 0.05) (Fig. 3). Similar to the mRNA transcription feature, the number of circRNAs decrease progressively, while the differential expression level expand along with the infection process (Fig. 3). In GO terms which circRNAs aligned to, several GO terms showed high frequency, for example: (1) ATP binding (GO:0005524), (2) catalytic activity (GO:0003824), (3) metabolic process (GO:0008152), and (4) intracellular (GO:0005622) (Fig. 6C).

**Figure 6.**
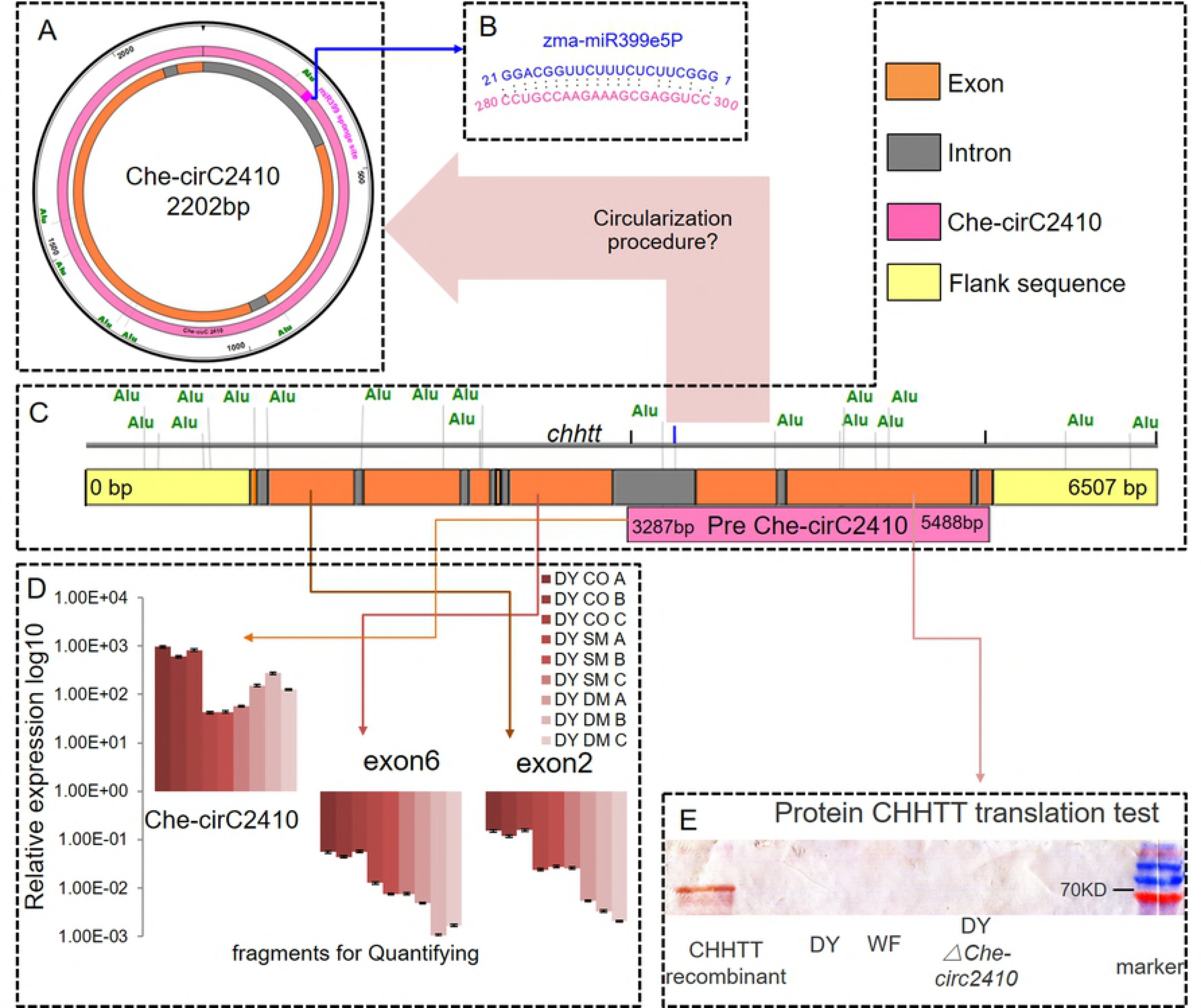
The circRNA differential status between high-virulence DY and low-virulence WF. (A) MA diagram for displaying the differentialcircRNA of each infection stage, the Y-axis represent the log2 fold change, the X-axis represent the gene mean normalized counts, the red spots mean that the differential is significant (P<0.05). (B) the length distribution of circRNA length. (C) the gene ontology(GO) distribution which circRNA fitted in.

### 4. Weighted gene coexpression network analysis (WGCNA) reveal virulence related genes

WGCNA is an useful tool for revealing the connection between co-expression transcripts and the virulence factors of microbe(40–42). In this, WGCNA analysis divided the race O transcriptome into 17 modules (Fig. 7A), and the correlations of each module with the disease IOD value was calculated and generated a heat map (Fig. 7B). Four modules had positive correlation with virulence, namely, the black (correlation: 0.74), brown (correlation: 0.91), cyan (correlation: 0.76), and midnight blue modules (correlation: 0.49). The function annotation revealed that most genes of the brown module converged on degrading enzymes, , the gene functions in the black module included transferase for nitrogenous groups, NADPH:quinone reductase, pyridoxal phosphate, GTPase, and ARF guanyl nucleotide exchange factor.

**Figure 7.**
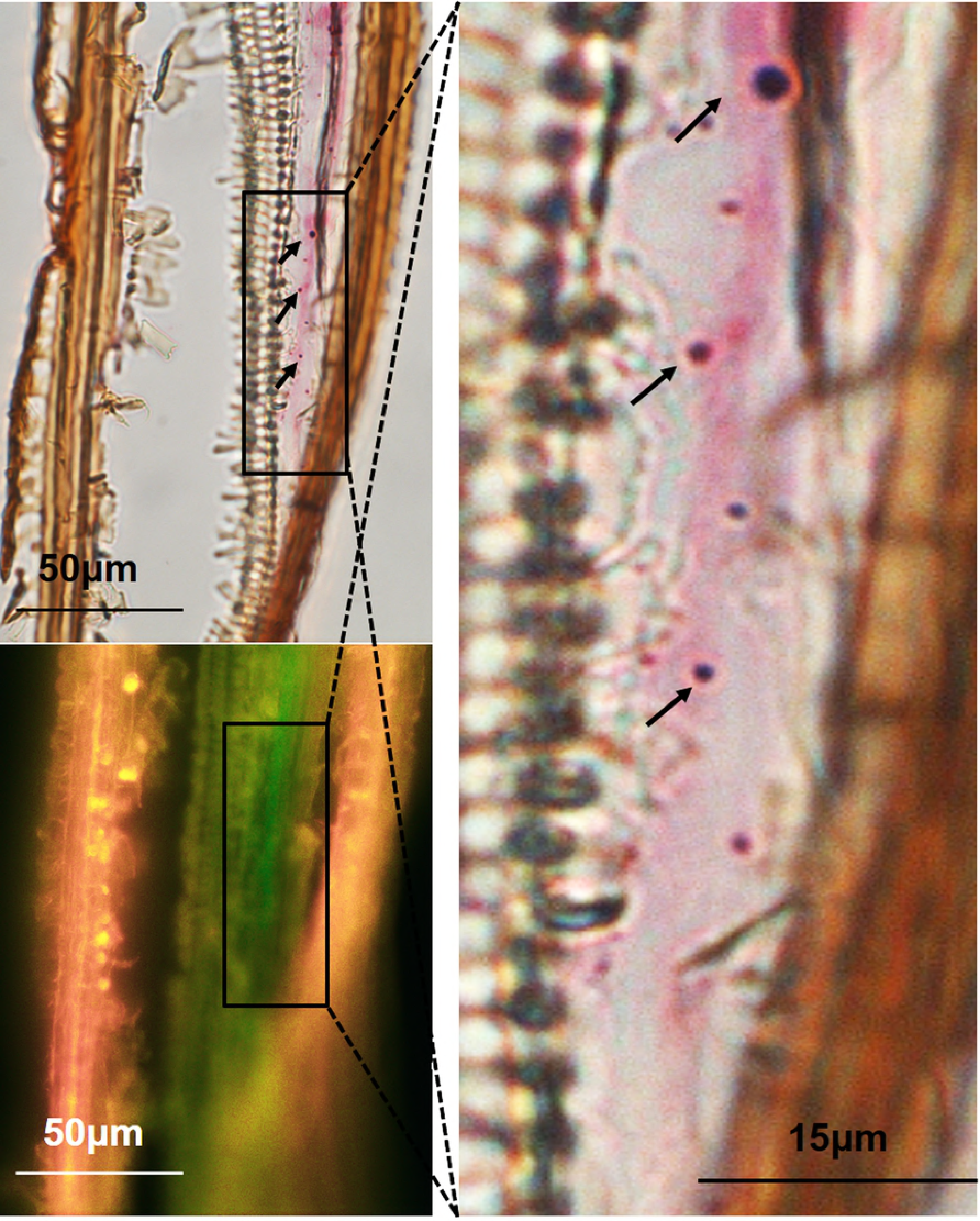
Modules of the mRNA, circRNA, and lncRNA transcripts revealed by WGCNA analysis. (A) Gene modules. The Y-axis represent the module name,the X-axis represent factors (strains and virulence) for correlation analysis, and correlation of each combination was listed in the module blocks. The scale of correlation is from the highest 1 (in color red) to the lowest −1 (in color green). (B) The midnight blue module demonstrated by heatmap.The mean expression level of each transcript was calculated using log2, and the expression scale ranges from 0 to 7. The X-axis represen each sample. The Y-axis represent the gene codes from 1 till 50, they are:1 Che-circ2410;2 Che-circ3417;3 Glycoside Hydrolase Family 28 protein;4 Ferric reductase;5 Mitochondrial/plastidial beta-ketoacyl-ACP reductase;6 Phosphotransferase;7 Nucleolar GTPase;8 Mitochondrial Fe2+ transporter;9 Sel1-like repeats;10 Cellobiose dehydorgenase;11 dTDP-4-dehydrorhamnose 3,5-epimerase;12 ER-Golgi vesicle-tethering protein p115;13 Rho GTPase-activating protein;14 membrane protein, contains DoH and Cytochrome b-561/ferric reductase;15 Karyopherin (importin) beta 1;16 Splicing coactivator SRm160/300;16 Splicing coactivator SRm160/300;17 Adenylosuccinate lyase;18 Exocyst protein Sec3;histone acetyltransferase;19 Oligosaccharyltransferase, gamma subunit;20 unknown;21 NmrA-like family;22 unknown;23 phosphoglycerate mutase;24 cyclin-dependent protein kinase regulator Pho80;25 Peptidase family S41;26 unknown;27 Peroxidase, family 2;28 unknown;29 40s ribosomal protein S27;30 isoamyl alcohol oxidase;31 60S ribosomal protein;32 nucleolar protein 5A;33 Sterol reductase/lamin B receptor;34 Uncharacterized integral membrane protein;35 Homeobox protein;36 Endoplasmic reticulum vesicle transporter;37 Dolichyl-phosphate-mannose;38 AMP-binding enzyme;39 3-oxoacyl CoA thiolase;40 Synaptic vesicle protein EHS-1;41 Homoaconitase;42 Alpha tubulin;43 Plasma membrane H+-transporting ATPase;44 Che-circ2414;45 Transcription factor TCF20;46 Endoplasmic reticulum vesicle transporter;47 26S proteasome regulatory complex;48 Myosin;49 Signal peptidase complex subunit;50 Vesicle coat complex COPII

Although the midnight blue module possessed a low correlation with virulence, its gene function was the most abundant: including polygalacturonase, phosphatidylinositol 3-kinase, acetyl-CoA C-acyltransferase, poly(A)-specific ribonuclease, protein transporter, 7S RNA binding, ferric-chelate reductase, triacylglycerol lipase, protein kinase CK2, 1-alkyl-2-acetylglycerophosphocholine esterase, cation transmembrane transporter, and transcription repressor. More important, in the midnight blue module, three circRNAs of over 10-fold differential level (Che-cirC2410, Che-circ3417, and Che-circ2414) between DY and WF implied that circRNAs may be biomarker of virulence variation (Fig. 7B). Sponging for miRNA is the primary function of circRNA (43–45), and the race O circular RNAs and microRNAs of *Zea mays* were performed the interaction prediction audaciously. A total of 19 interaction matrices were generated (S1_dataset), in which at least one pathogen circRNA target on at least one miRNA of host. The predicted miRNA families include MIR396, MIR159, MIR167, MIR168, MIR172, MIR395, MIR398, MIR399, MIR408, MIR482, MIR2118, and MIR2275. Some noncoding RNA interaction matrices are simple, such as: Che-cirC2410 was predicted to interact with the host zma-miR399e-5P (Fig 8, Fig 10A, Fig. 10B), and zma-miR399e-5P was predicted to target 35 maize transcripts (Fig. 8), and diverse significant genes, such as serine/threonine kinases, RING finger were found among these targets (Table 2). However, some interaction matrices are complicated: two pathogen circRNAs (circ3462, circ3475) were predicted the interaction with four host miRNA, and the excess targets of miRNA beyond our cognitive (S1_dataset, page1).

**Table 2.**
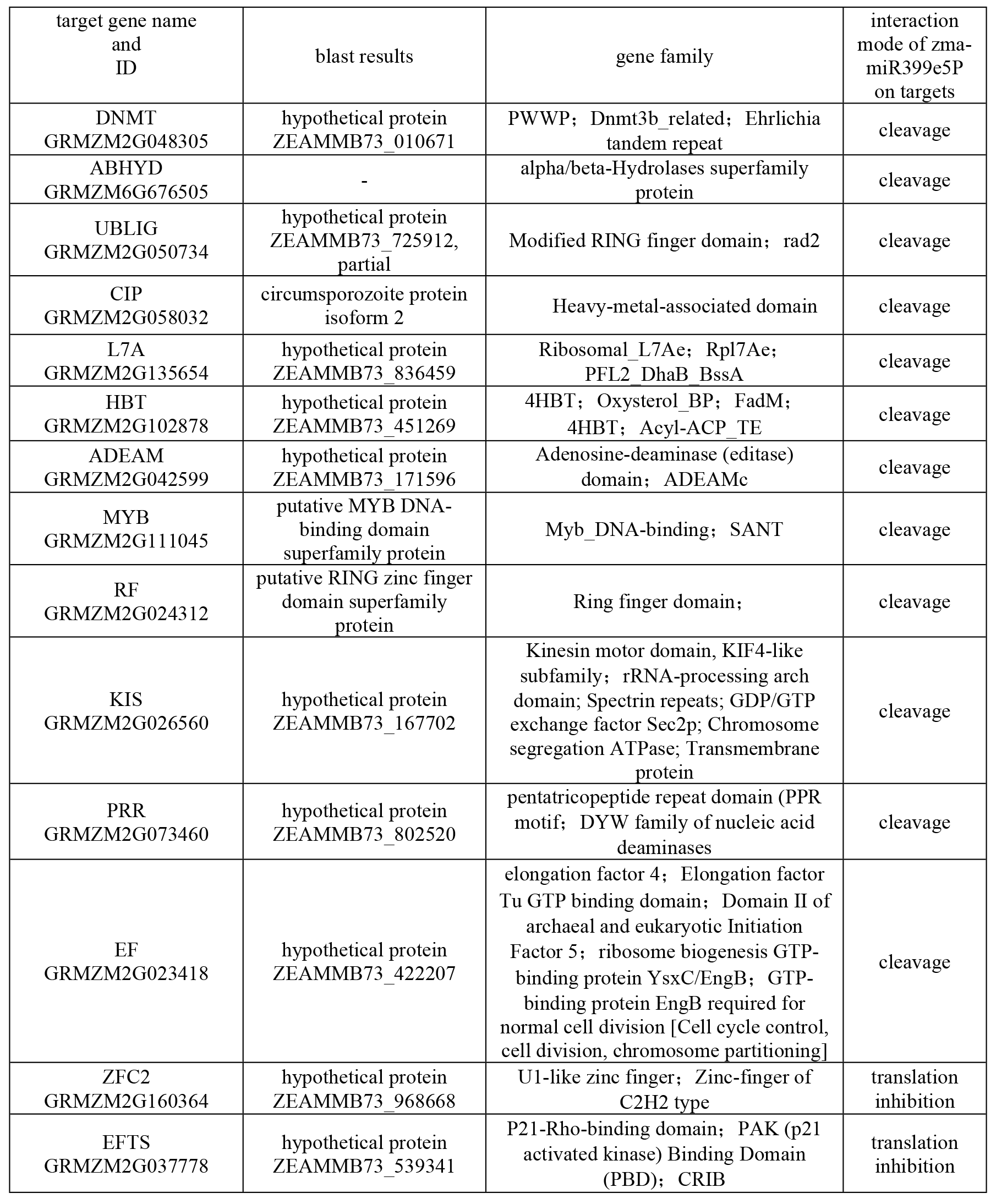
Targets gene of zma-miR399e5P and functional annotation

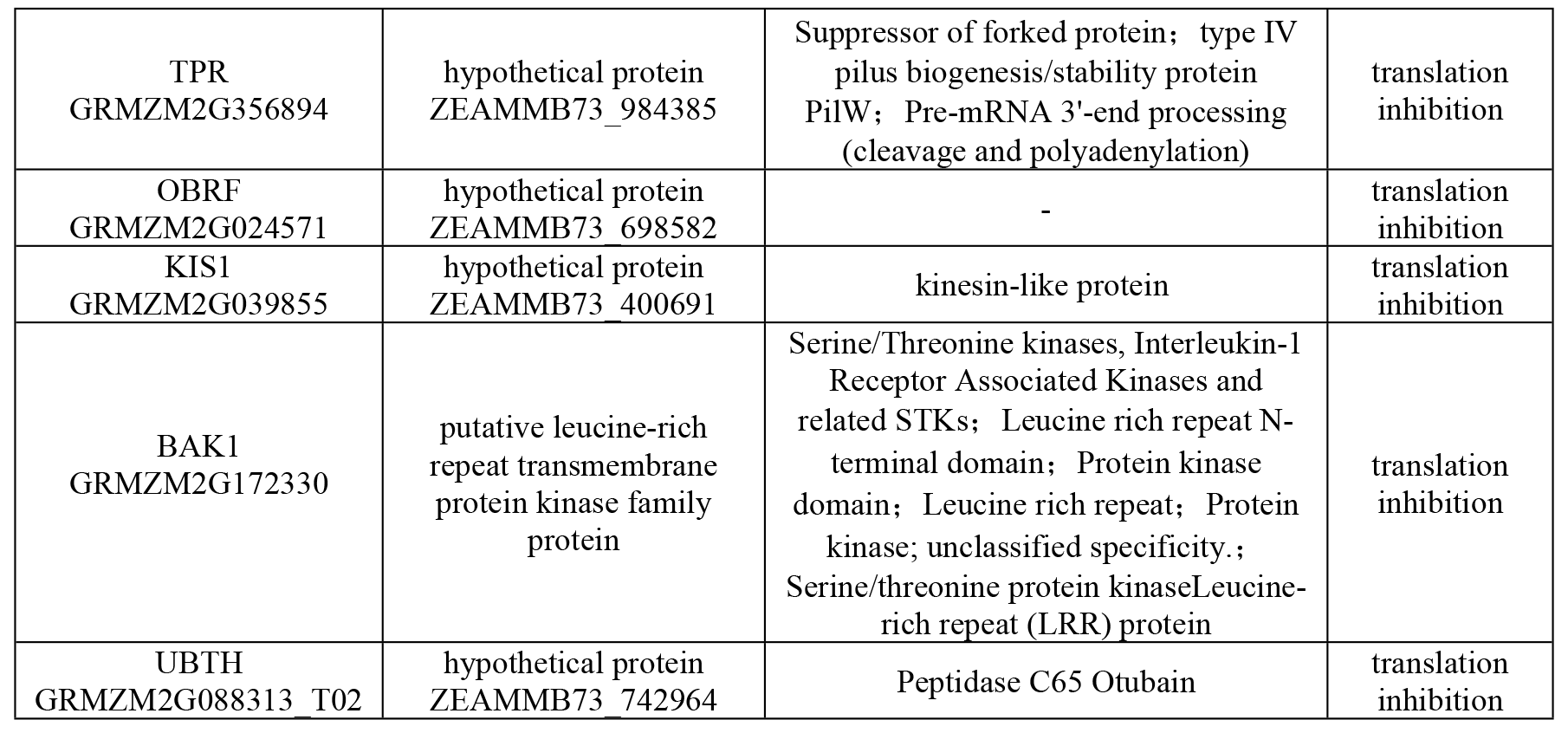

**Figure 8.**
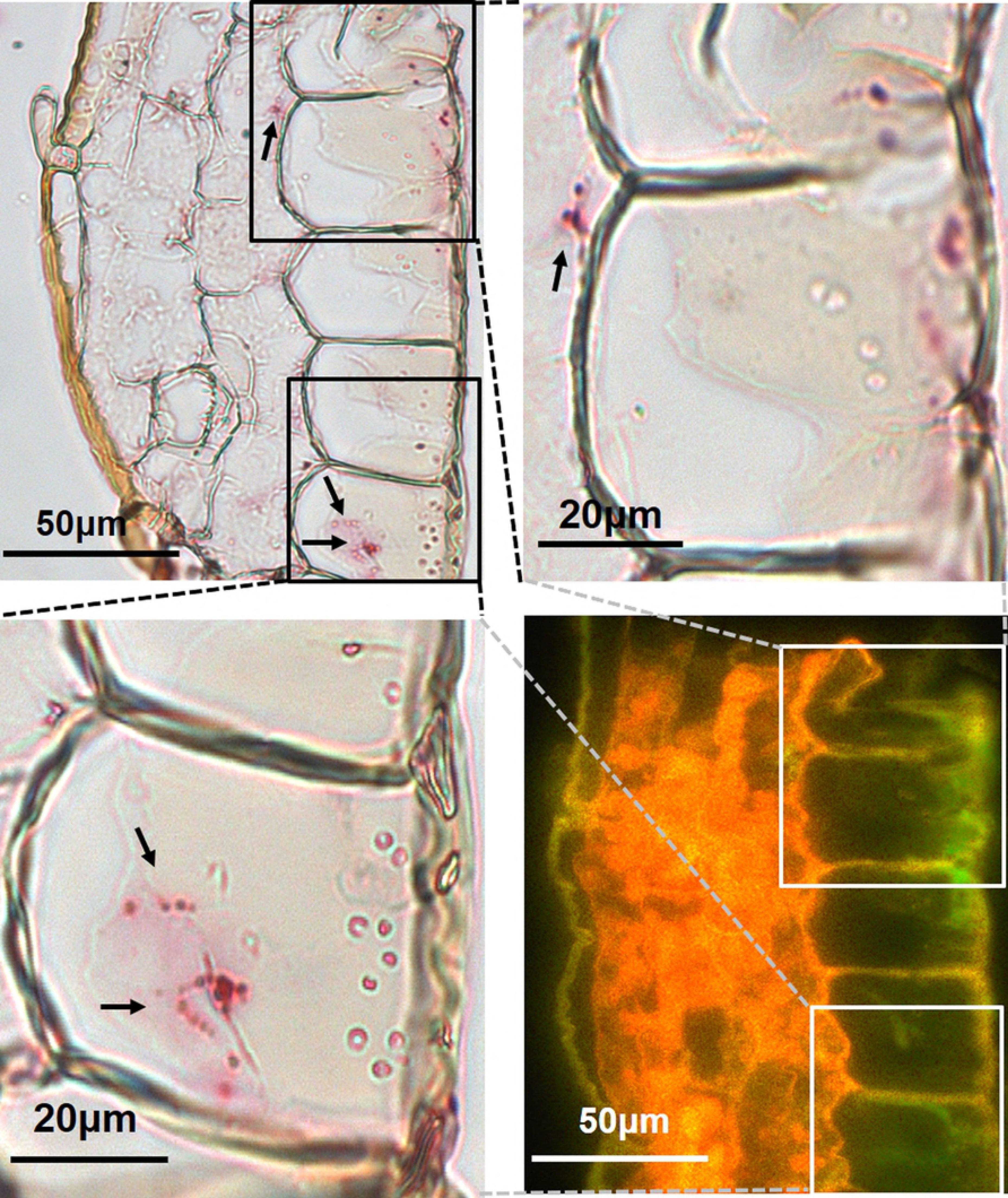
Pathway from the *C. heterostrophus* race O circRNA Che-circ2410 to the host B73 zma-miR399e5P and the predicted targets of miRNA. circRNAs are indicated by cyan diamonds, miRNAs are indicated by red spheres, and the mRNA targets are indicated by green arrowheads.

### 5. Molecular characteristics and functions of noncoding RNAs related to *C. heterostrophus*-host maize interaction

Insensitivity to the digestion of Rnase R (3′-5′ exoribonuclease) is one of the circRNA characteristics(45). After treatment of RNase R, the relative expression level of Che-circ2410 increased for 4.7 fold, by contrast, relative expression level of the linear RNA ACTIN1 decreased for 5 fold (Fig. 9B). The predicted length of Che-circ2410 is 2202bp, the fragment amplified by using divergent primers is around 190bp, and the fragment amplified by convergent primers is around 2100bp, the total length of two fragments are according to the Che-circ2410 (Fig. 9A). Hence, evidence above tells that Che-circ2410 is a naturally true circRNA of *C.heterostrophus* race O.

**Figure 9.**
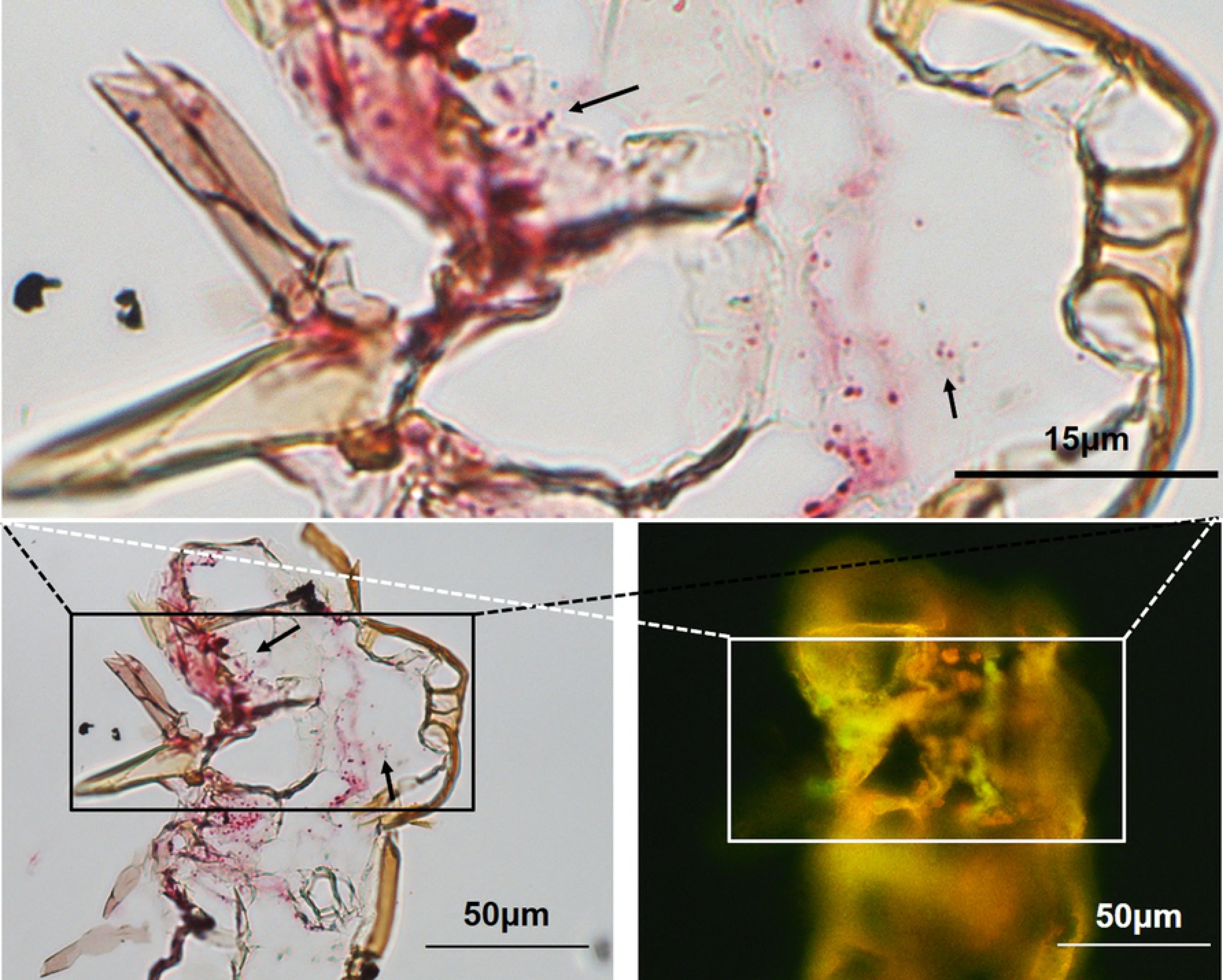
The characteristics of Che-circ2410. (A) The convergent and divergent fragments on agrose gel. The convergent and divergent fragmen add up to 2200bp, which accord with the predicted length of the Che-circ2410. The white sket map inside A exhibit the primer designing strategy for circRNA verification. (B) Survivability of Che-cirC2410 to RNase R. A total of 10 μg total RNA from the DY colony mycelium was digested by RNase R for 1 h. The concentration after digestion was 36 ng, demonstrating that most linear RNA was erased. A total of 500 ng digested RNA and 500 ng undigested RNA were reverse transcribed, and then the expression levels were quantified using real-time PCR. The actin content of the undigested sample was introduced as the calibration sample. All experiments were performed thrice.

**Figure 10.**
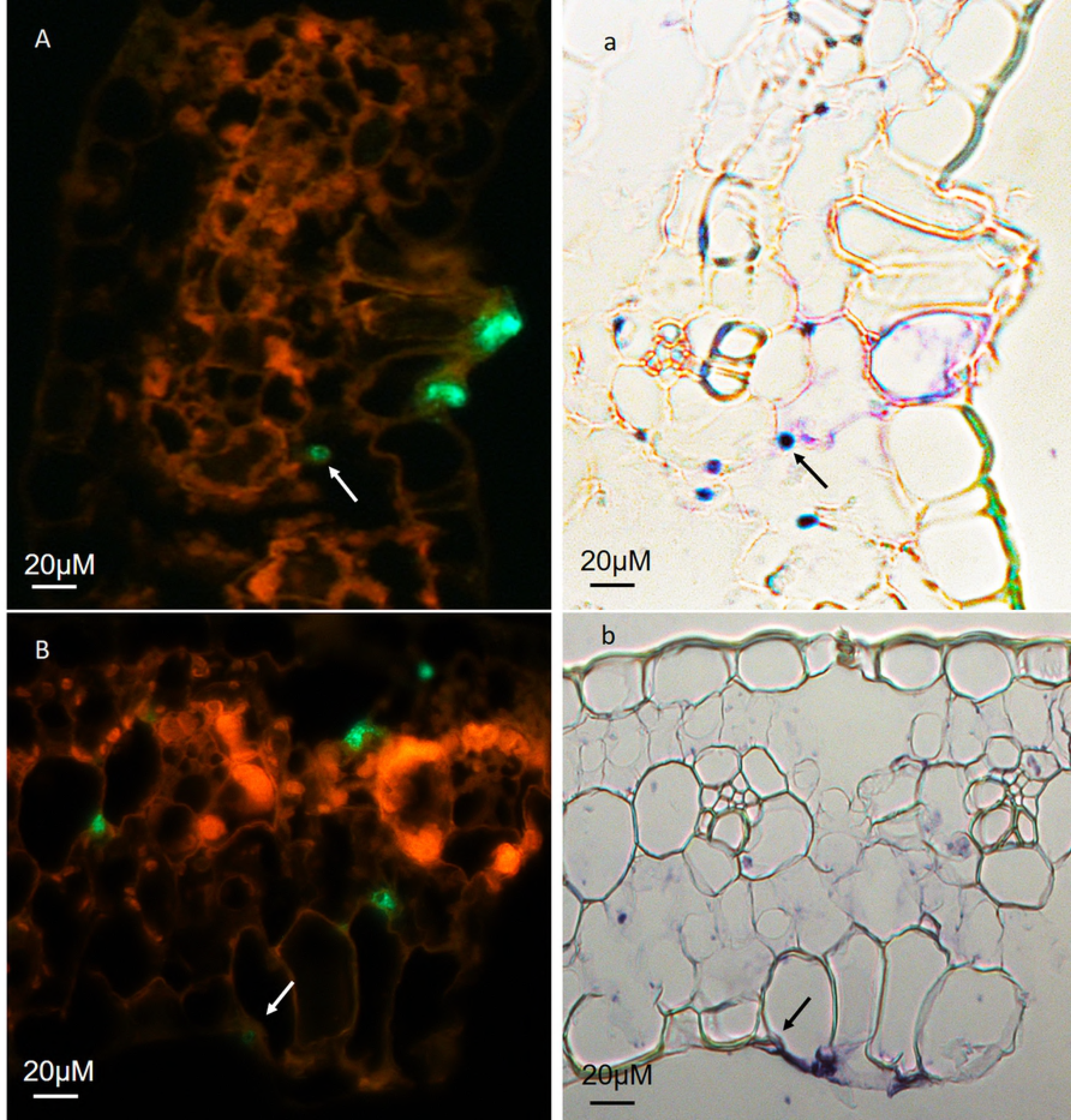
Anatomy,exon expressin and protein translation test of the *chhtt* gene and Che-circ2410. (A) The length of Che-cirC2410 is 2202 bp, and it is located at the 3287-5488 bp of *chhtt*. (B) The site from 280 bp to 300 bp of Che-cirC2410 is the sponge site for zma-miR399e5P. The maximum energy to unpair the Che-cirC2410-zma-miR399e5P target site (UPE) is 15.283. (C) The length of the *chhtt* gene is 6507 bp (including 1000 bp upstream flank and 1000 bp downstream flank sequence). A transcriptional level mutation (from GTC to CAG) was observed at the exact junction site. The feature sequence prediction demonstrated the presence of 9 exons, 8 introns, and 18 Alu features (nucleotides: AGCT).One pair of homologous sequence (located at 574-584 bp and 6251-6261 bp) and one pair of reverse complement sequence (located at 26712681 and 5974-5964 bp) were found.The orange bar represents the exons, the grey bar represents the introns, the rose-red bar represents the circRNA Che-cirC2410, and the yellow bar represents the flank sequence. (D) Three RNA fragments were tested to obtain the expression levels. The exon2 and exon6 comprised nonhomologous sequences of Che-cirC2410. The junction sequence was tested at the expression level by using a pair of diverge primers. The relative expression was log10 calculated, the circRNA expression was extremely high, by contrast, the exon2 and exon6 expression were almost zero. (E) The ability of protein translation was tested by a monoclonal antibody that targets the peptide (translated by the ninth exon),the CHHTT recombinant was used for inducing antibody and positive control of western-blot, there is a clear band on the nitrocellulose membrane, however, there was not antigen detected from the nature protein extracted of strain DY, WF and *DY Che-circ2410*. All experiments were performed thrice.

Data which is helpful for understanding the production mechanism of fungi circRNA was analyzed: Che-circ2410 is an exon-intron type circRNA, which is made up of 3 exons and 3 introns (Fig. 10A), the pre RNA of Che-circ2410 is from 3287bp till 5488bp of *chhtt*, a transcriptional level mutation was observed in Che-cirC2410: GTC mutated into CAG at the exact junction site (gene *chhtt* position at 5486 bp) (Fig.10C). In the Che-circ2410 original gene *chhtt*, 18 Alu(46) features from upstream flank(1 KB) till downstream flank(1 KB) were found (Fig. 10C). There is a pair of reverse complement sequence at the flank of *chhtt*, and a pair of homogeneous sequence at the flank of Che-circ2410 (Fig. 10C). To date, the linkage between the transcriptional level mutation and circRNA production remains unknown. However, Chen, et al. found that the exon circularization was mediated by complementary sequence, and the Alu sequence was involved in the alternative circularization(47–49). Beside the Che-circ2410, there were other 4 isoforms of circRNA produced from the single gene *chhtt*, namely circRNA2414(924bp), circRNA2415(467bp), circRNA2416(443bp), circRNA2417(327bp), however, the isoforms lack the sponge site for zma-miR399e5P, which imply insignificance for the pathogen-host interaction.

Two fragments of the mRNA which shared no intersection with the Che-circ2410 and one circRNA junction fragment were detected the expression simultaneously. The quantitative PCR showed that the transcription of mRNA was almost zero(Fig. 10D). The highest and lowest relative expression of the junction fragment were 964.3 at the CO stage and 43.51 at the SM stage, respectively (Fig. 10D). Western blot analysis revealed that no positive band was detected by the monoclonal antibody (Fig. 10E), *chhtt* is recognized as a pseudogene that only produces RNAs.

Logically, the meaning of biomolecular limiting inside or transfering outside the pathogenic fungi cytomembrane varies considerably, only if the Che-circ2410 moves into the host tissue, it has chance to sponge miRNA. For this, the subcellular location of Che-cirC2410 and zma-miR399e5P in the interaction tissue was exhibited by using BaseScope™ in situ hybridization (ISH)(50–53) and locked nucleic acid (LNA) ISH(54–56), respectively. Inspiringly, much Che-cirC2410 was detected by circRNA probe in the diseased host tissue, the Che-circ2410 not only gathered around infecting mycelium (Fig. 11), and also disperse in the motor cells (Fig. 12), mesophyll cells (Fig. 13). Additionally, the Che-circ2410 was not found in vascular bundle. Considerable zma-miR399e5P was detected in the cytoplasm of the mesophyll cells but emerged rarely in vascular cells (S3 Fig.), when the leaf was infected with the pathogen, the density of the zma-miR399e-5P signal declined in mesophyll cells (Fig. 14

**Figure 11.**
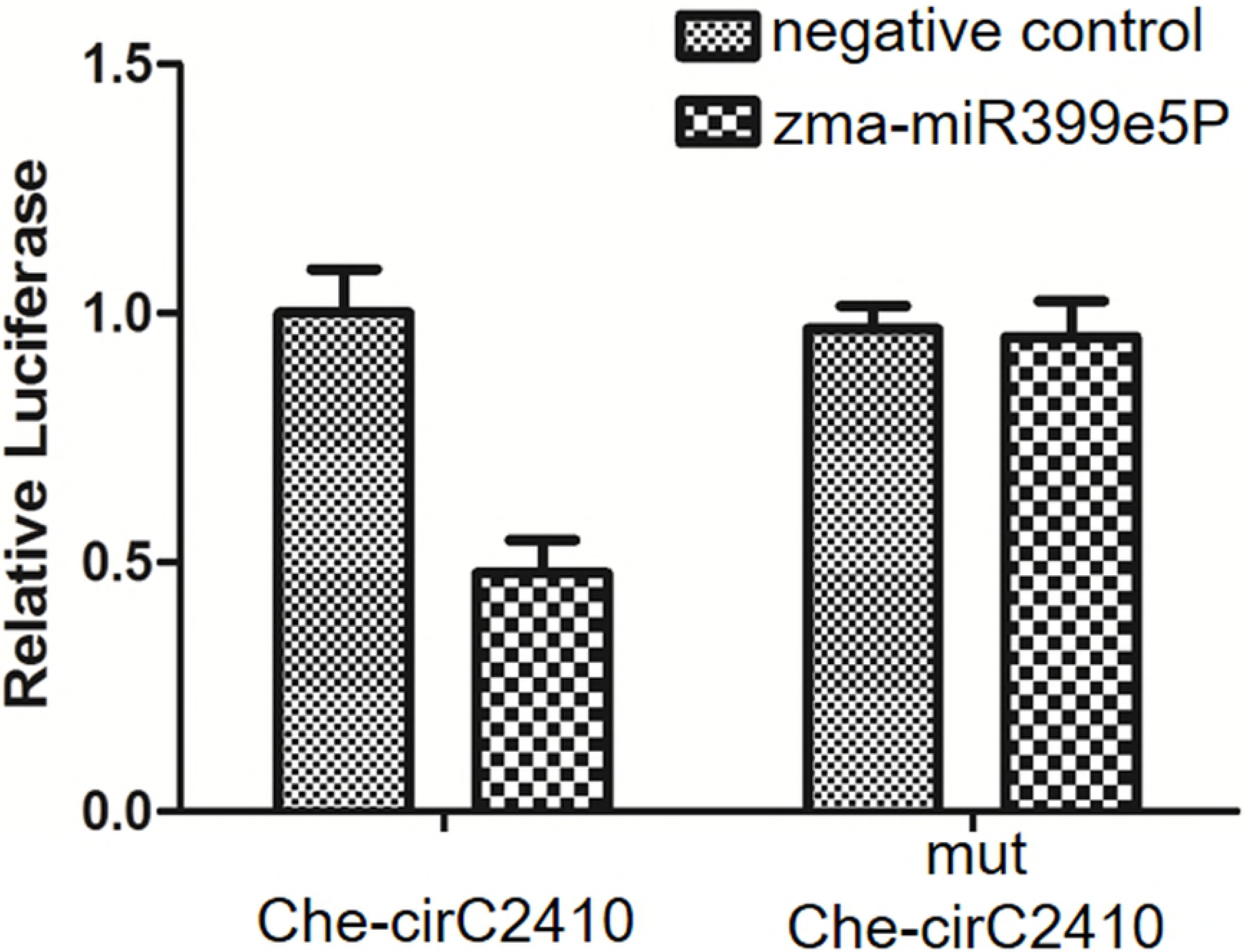
Subcellular location of Che-cirC2410 around the *C. heterostrophus* race O mycelium. Firstly, the infected leaf was fixed in paraffin and sliced along the vein. The infecting pathogen DY in the surrounding veins was located under a fluorescence microscope (bottom-left photograph; excitation wavelength: 485-560 nm; emission wavelength: 520-610 nm). After the BaseScope^TM^ in situ hybridization workflow, the Che-cirC2410 was located under a bright-field microscope (top-left and right amplified photographs). The BaseScope^TM^ signal spots were mostly observed on the outline of the mycelium, and signal spots were also found outside of the mycelium. Scale bars: 50 and 15 μm.

**Figure 12.**
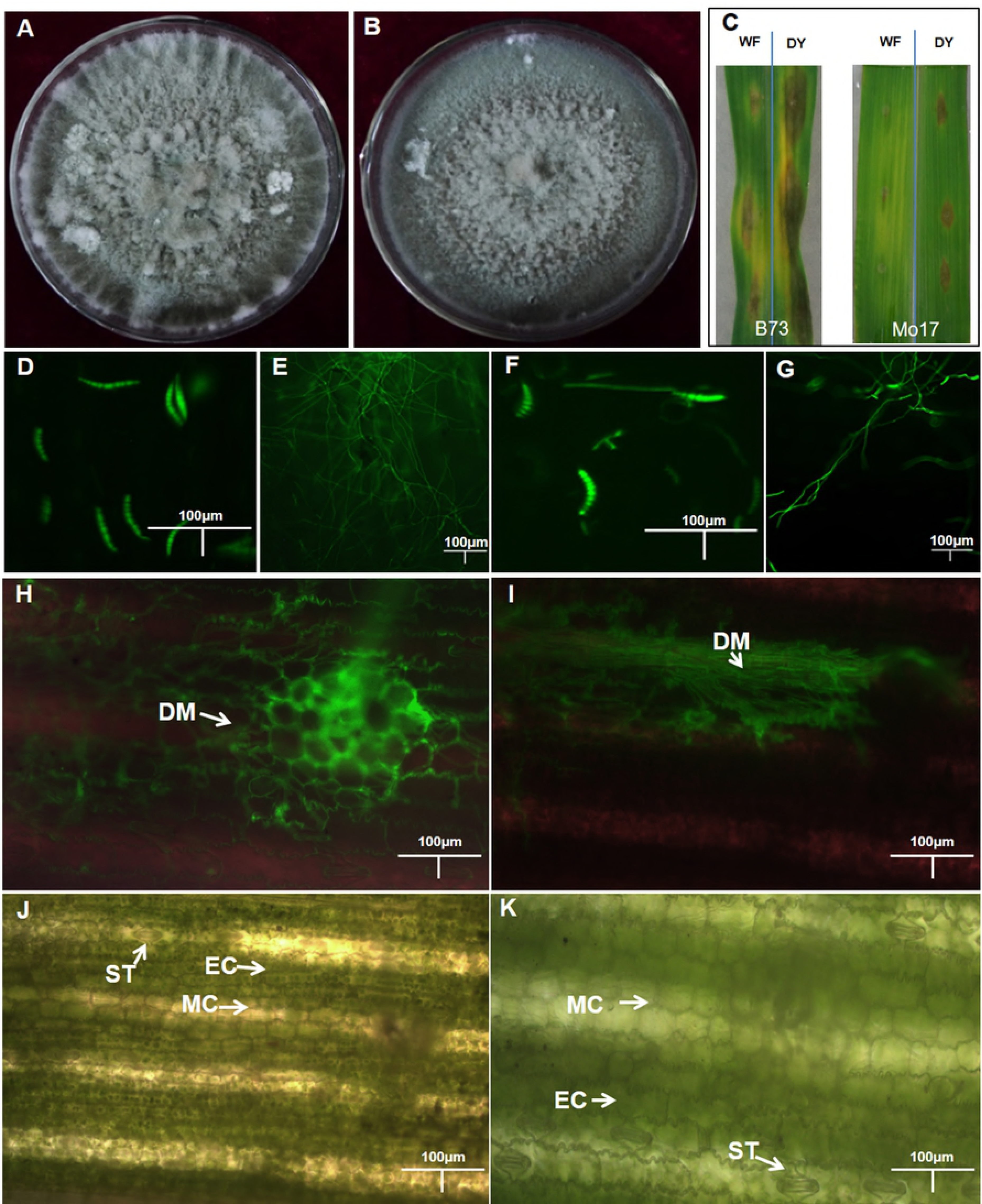
Subcellular location of Che-cirC2410 in the host motor cell. The workflow was described as same as in the caption of Fig11. Scale bars: 50 and 15 μm.

**Figure 13.**
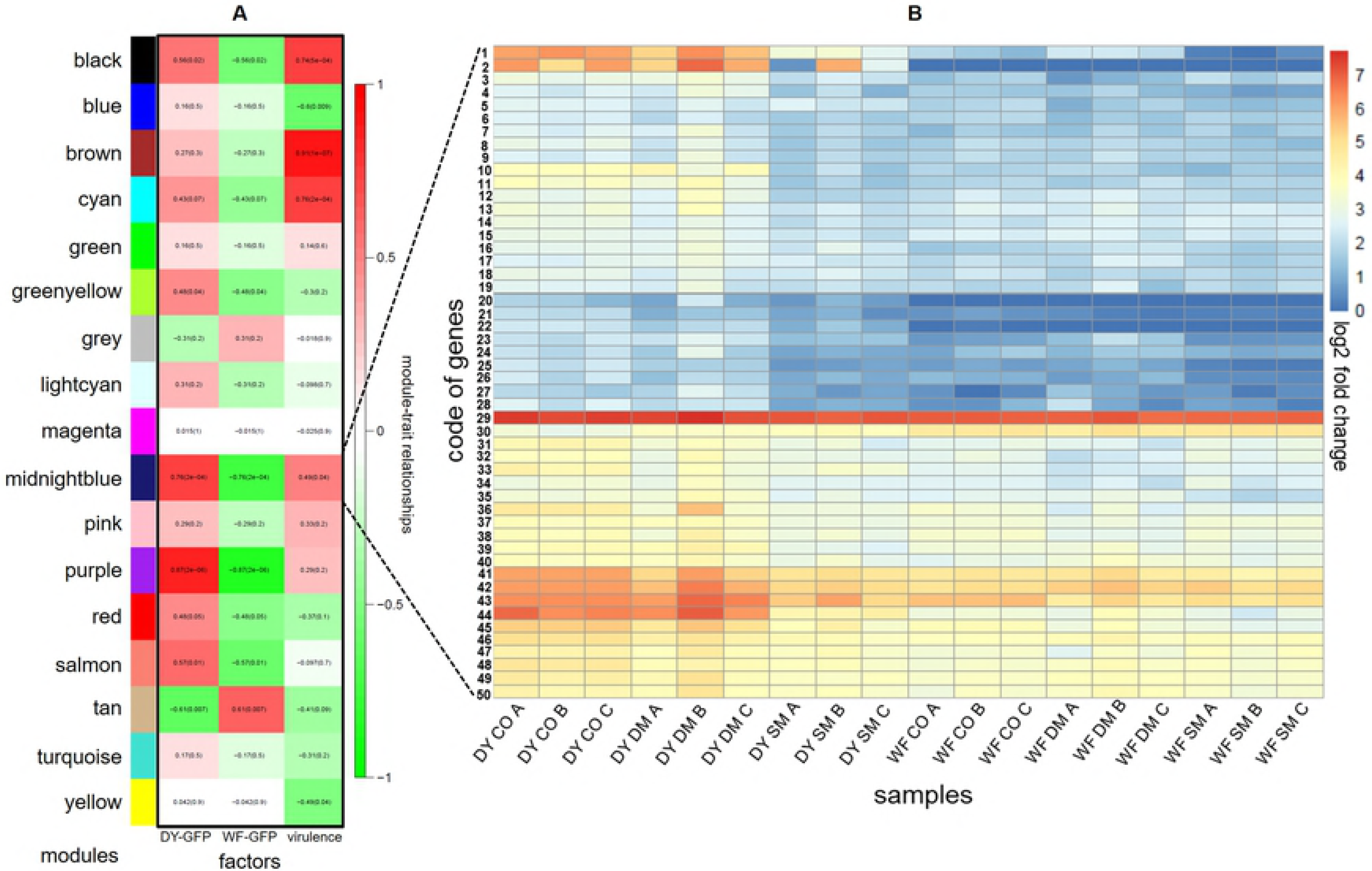
Subcellular location of Che-cirC2410 in the host mesophyll cell. The workflow was described as same as in the caption of Fig11. Scale bars: 50 and 15 μm.

**Figure 14.**
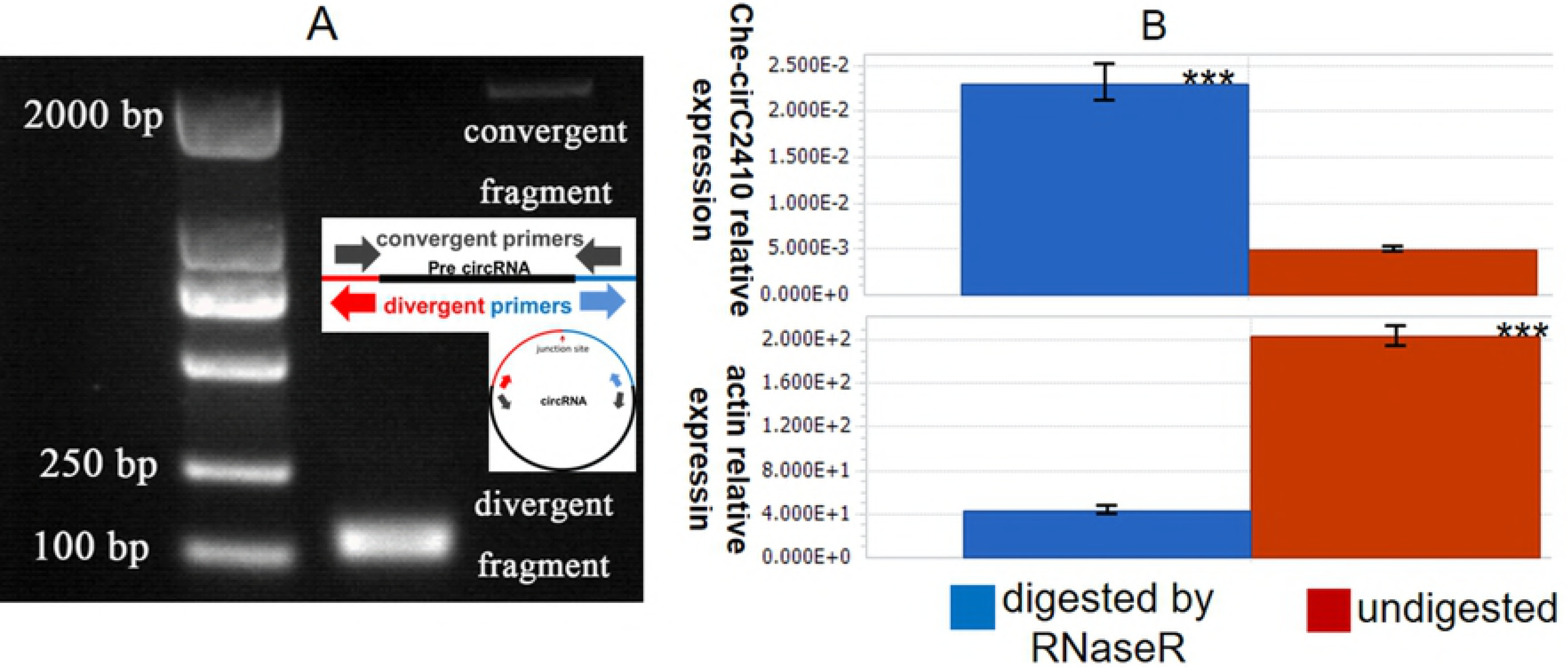
Subcellular location of zma-miR399e5P in the *C. heterostrophus*-B73 infection complex. First, the infected leaf was fixed in paraffin and sliced perpendicular to the vein. The infecting pathogen DY was located in the motor and mesophyll cells under a fluorescence microscope (left photograph, excitation wavelength: 485-560 nm; emission wavelength: 520-610 nm). After the in situ hybridization by using a locked nucleic acid (LNA) probe, the zma-miR399e5P was located under a bright-field microscope (right photograph). The LNA probe signal spots were found in mesophyll cells with a low concentration. On the first tissue at the top-right photograph, 15 zma-miR399e5P spots were observed. On the second tissue at the bottom-right photograph, rare spots were observed. Scale bar: 20 μm. All experiments were performed thrice.

To further determine the sources of the two noncoding RNAs, we tested the pure plant and fungal colony. The zma-miR399e-5P were detected in maize only, while Che-cirC2410 was detected in the fungal pathogen only (S3 Fig., S4 Fig.).

The ISH results provide the probability of interface between Che-circ2410 and zma-miR399e5P, additionally, a dual-luciferase reporter system was used to test this (Fig. 15). The fluorescence intensity of the negative control was 10.266, and that of the experimental group, Che-cirC2410-zma-miR399e5P, was 48% of the negative control (4.907, P = 0.0012) (S4_Table). The results indicated a significant variation resulting from the sponge. Moreover, to exclude the false positive that the reduction of fluorescence intensity generated from a potential plasmid sequence, we introduced a nucleotide mutation of the Che-cirC2410, and the fluorescence intensity was recovered to 95% of the negative control (Fig. 15).

**Figure 15.**
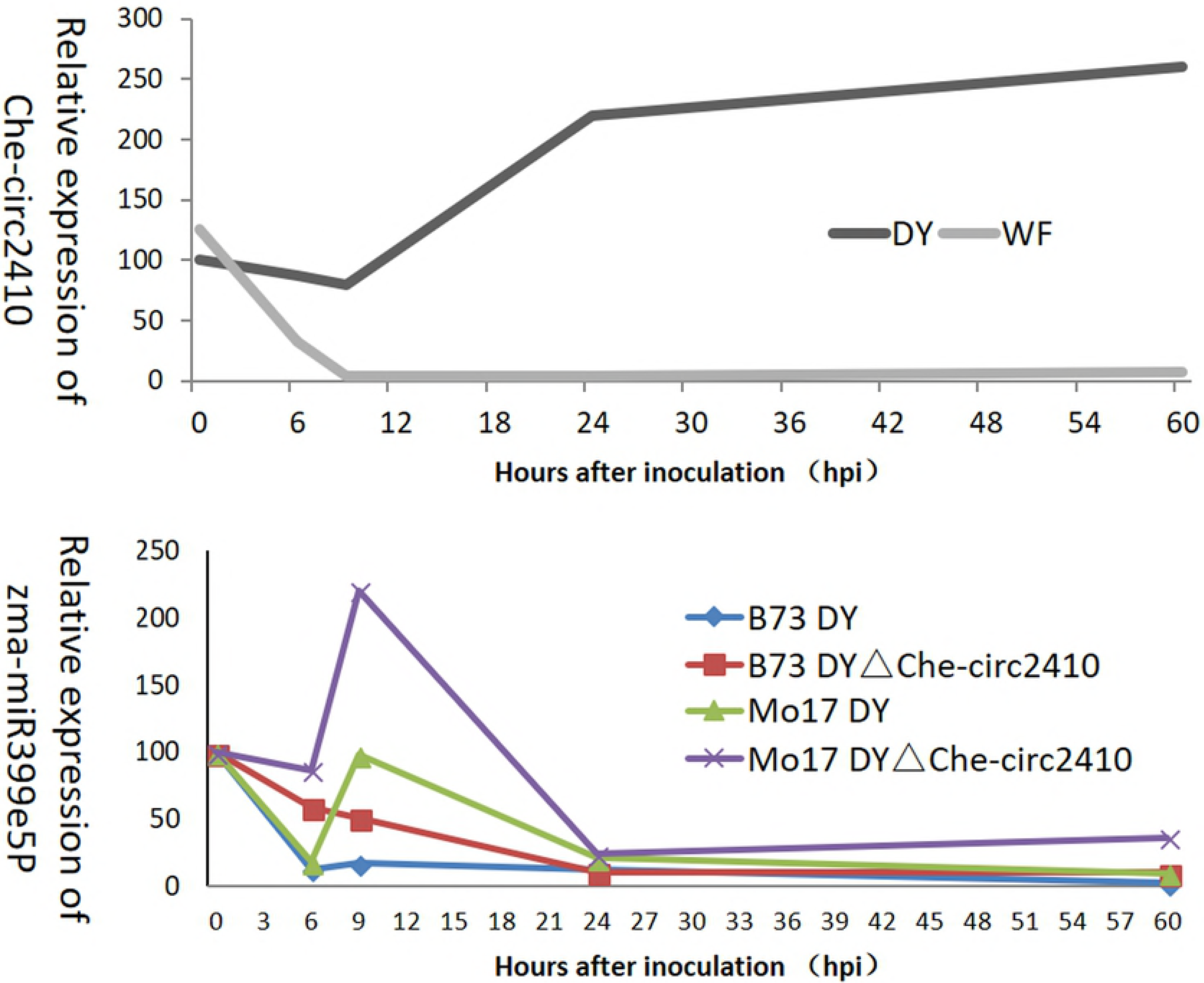
Dual-luciferase reporter system to test the interaction between Che-cirC2410 and zma-miR399e5P. The relative luciferase value of the experimental group was 4.9, which exhibited a significant reduction compared with the negative control value of 10.266 (P = 0.0012). After the sponge site of Che-cirC2410 was mutated to nucleotide, no relative luciferase variation was observed between the experimental group and negative control. The significant difference and nucleotide mutation settings demonstrate the interaction between zma-miR399e5P and Che-circ2410. All experiments were performed thrice.

The interaction between noncoding RNAs was demonstrated in vitro, more, the expression of zma-miR399e5P was evaluated in vivo. When the Che-cirC2410 of *C. heterostrophus* was efficient, the expression of zma-miR399e-5P in the infected B73, Mo17 inbred lines declined sharply at 12 hpi, and the expression level of miRNA of Mo17 slightly increased at 24 hpi. The zma-miR399e5P relative expression level was 2.5 in B73 and 9.58 in Mo17 at 60 hpi. When the Che-circ2410 of race O was deficient, the expression level of zma-miR399e-5P was higher than that in B73 infected by DY*△ Che-circ2410* at 12, 24, and 60 hpi and in Mo17 at all infection periods (Fig. 16). The average expression level of zma-miR399e-5P was 29.1 in B73 infected by DY, and 46.2 in B73 infected by DY*△ Che-circ2410*. The average expression level of zma-miR399e-5P was 49.4 in Mo17 infected by DY, and 93.5 in Mo17 infected by DY*△ Che-circ2410*. The Che-cirC2410 knockout mutant induced a higher zma-miR399e-5P latitude in the hosts infected by DY*△ Che-circ2410* than in those infected by the pathogens with efficient Che-cirC2410. The expression data was according to the ISH of zma-miR399e5P.

**Figure 16.**
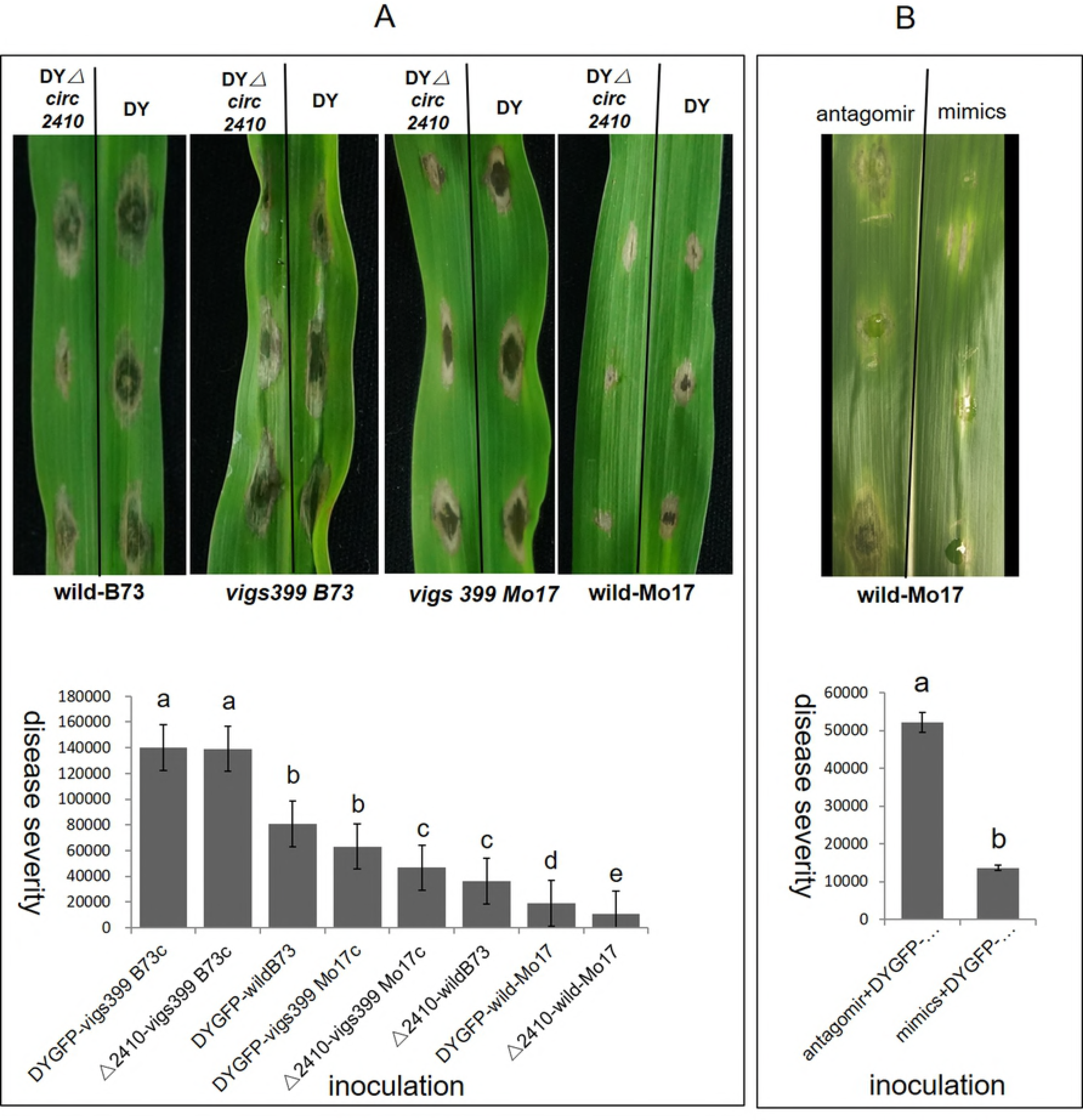
Expressions of Che-circ2410 and zma-miR399e5P in the infection process. (A) The expression of Che-circ2410 in strain DY and WF. (B) The expression of zma-miR399e5P in inbre lines B73 and Mo17 infected by DY and *DY△ Che-circ2410*. The healthy plants at 0 h after inoculation (hpi) were used as mock samples.

What are the function of two novel noncoding RNA in phytopathology? For answering, the pathogen Che-circ2410 knockout mutant (DY*△ Che-circ2410*) and host zma-miR399e5P silencing plant(*vigs399Mo17*, *vigs399B73*) were built (S3 Fig., S5 Fig.). The DY*△ Che-circ2410* strain exhibited a 0.57-fold (infecting wild-type Mo17) and 0.42fold (infecting wild-type B73) reduction in virulence compared with the Che-cirC2410-efficient strain DY. The disease severity was 3.29-fold higher in *vigs399Mo17* (infected by DY) than in wild-type Mo17 (infected by DY) and 1.73-fold higher in *vigs399B73* (infected by DY) than in wild-type B73 (infected by DY) (Fig. 17A). Thus, the positivities of Che-circ2410 for virulence and zma-miR399e5P for resistance were revealed, however, for now, we lack the methods for overexpressing fungi circRNA or maize miRNA in vivo. Alternatively, the antagomir and mimics of zma-miR399e5P were synthesised and injected into host for substituting the Che-circ2410 sponge region and zma-miR399e5P separately, and the disease severity of antagomir-treatment is 3.84-fold higher than the mimics-treatment (Fig. 17B).

**Figure 17.**
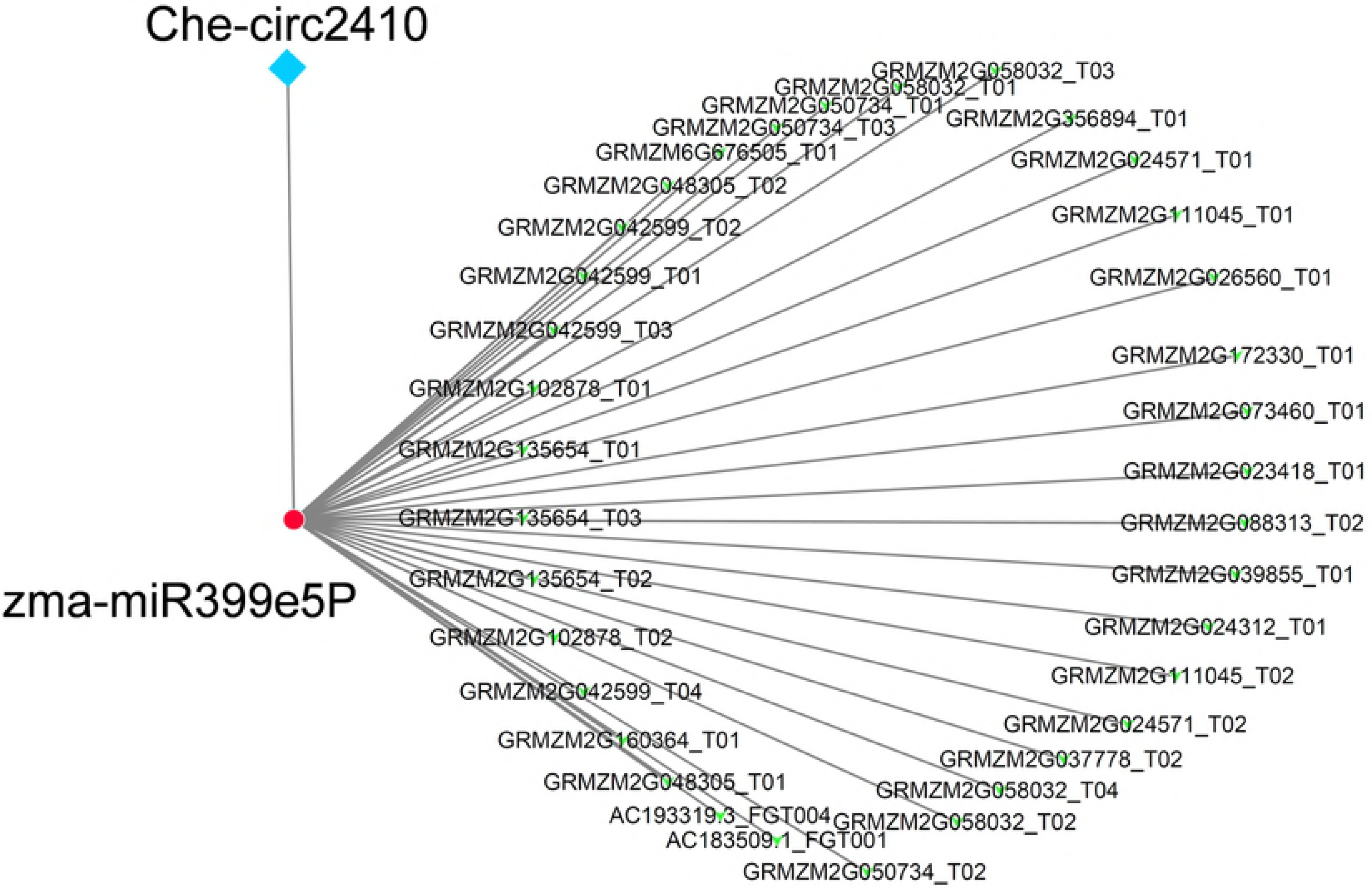
Disease severity of the different infection treatments,which demonstrated the important contribution of zma-miR399e5P in host resistance and Che-circ2410 in pathogen virulence. (A) The bottom labels of disease photograph are the host treatments by VIGS and wild controls. The top labels are the pathogen treatments by DY and *DY△Che-circ2410*. The wild pathogen and mutation were inoculated on same foliage at two sides of the vein (the veins were delineated by the black line). The disease severity was calculated and the significance of difference was labeled by alphabet. (B) For simulating the over-expression, the leaves in vitro were injected the mimics and antagomir of zma-miR399e5P. The differential disease severity was exhibited on the photography and the significance of difference was calculated and labeled by alphabet.

## Discussion

### 1. The infection of *C.heterostrophus* race O

The dense bundles of thick mycelium (DM) was firstly reported in *C.heterostrophus* race T by Horwitz(57), and the location of race T DM was uncertain, by contrast, the race O DM was found with high tropism to the motor cell of host. Motor cells/bulliform cells are large and short of cell inclusion, and related to the leaf rolling under biotic/abiotic stress (58). There are cross epidermal cells above the motor cell, which protect leaves from the fungi infection(59–61). However, in the race O-maize interaction, the epidermal cells failed in rejecting the invading of pathogen, and exhibit tropism to the DM structure on the contrary. Hence, taking the epidermal cell as the SCLB resistance marker is open to discussion.

### 2. The methodology of whole transcriptome

This report is the one of the few presentation on the whole transcriptome (including mRNA, lncRNA, and circRNA) of microbes (62–64). In the classical RNA-seq work after single-cell level LCM, the global expression of mRNA by using oligodT primers is generally applied (11, 16, 17), while, for revealing the whole transcriptome, we used the DNA/RNA chimeric primer mix and SPIA^®^ Amplification (14, 65), and successfully acquired the mRNA, lncRNA and circRNA. However, it is imperfect that the miRNA of *C.heterostrophus* race O could not be sequenced in an one-off workflow. Considering the importance of miRNA in the host-pathogen interaction(66–69), it is necessary for revealing the miRNA world of *C.heterostrophus* in infection.

### 3. The mRNA transcription of *C.heterostrophus* race O in infection

The pathogenic significance of FTFD, TCDB, P450, SSP, NPS and PKS is undoubted(70–75), and all kinds of virulence related genes above were detected. Among the virulence related gene catalogs, SSP is most abundant in both coexpression and differential-expression analysis. PKS is the fatal virulence factor of *C.heterostrophus* race T, however in race O, most PKS genes down-regulated, only PKS9 and PKS15 up-regulated along with infection progress. The transcription of PKS in virulence differential race O strains were unequal, and indicated that PKS not the lethal factor. The feature genes in each infection stage implied the significant genes activity beside virulence related genes, and provided another potential targets for designing fungicide.

### 4. The noncoding RNA of *C.heterostrophus*

The number of circRNAs(2279) were 18-fold higher than the number of lncRNAs(169), Yuan et al. reported that circRNA differentially expressed in conidia and mycelium of *Magnaporthe oryzae*, and the circRNA number (8848) is more than *C.heterostrophus* race O (64). In fungi, circRNA was firstly reported in yeast (76, 77), and emerge in multiple pathogen fungi (62, 78), which implied that circRNAs may play important roles in pathogen infection. The focus of Che-cirC2410 was on account of 3 reasons: 1 Che-cirC2410 exhibited differential expression in two race O strains; 2 Che-cirC2410 was predicted a positive correlation with virulence; 3 Che-cirC2410 was predicted sponge interaction with zma-miR399e5P. MiRNAs, particularly the plant miR399 family, play critical roles in mediating plant nutrients balance and reaction to stresses. In the rapeseed, pumpkin and arabidopsis, miR399 was proved a signal for regulating the phosphate homeostasis(79–81). In *arabidopsis thaliana*, the expression of miR399f was regulated by a salt and drought stress responses protein AtMYB2, and miR399f was also found participates in responses to abscisic acid(82, 83).

In soybean, it was reported that nine miRNAs were regulated by noncoding endogenous target mimics (eTMs), and these noncoding RNA interactions were involved in lipid metabolism(84). *IPS1* is another eTM of *Arabidopsis thaliana*, it was reported an interaction with miR399, and the canonical target PHO2 of miR399 was interfered(85). In animals, the interaction among noncoding RNAs was described by the concept of competing endogenous RNA (ceRNA), which ideally connects split transcripts, such as lncRNA, circRNA, siRNA, miRNA, and mRNA(86, 87). In ceRNA, miRNA serves as the core bridge, and the targets for miRNA share the same miRNA response elements (MRE), however, the ceRNA and eTMS-miRNA are limited in single species. Cross-kingdom transporting and contact are the preconditions of effectors` functioning on host target, for instance, type III secretion system is a powerful weapon that pathogenic bacteria applied for delivering protein effectors into host cell (88, 89), effectors from oomycetes was delivered into soybean to hijack the host target(90). Even for effectors that are extracellular (such as Pep1, Pit2, Avr2), there is apoplastic space between the fungal hypha and the plant plasma membrane for effector-target meeting(74). Here, inspired by the ceRNA and eTMs hypothesis, we boldly speculate the cross-kingdom delivery of *C.heterostrophus* circRNAs, and target on potential host microRNA afterwards. The circRNA Che-cirC2410 was secreted into the host maize cells and sponged microRNA target zma-mir399e5P, and it is worthy for exploring other pathogenic noncoding RNA-host microRNA interaction events.

For now, we are still blind of two points: (1) the channel transporting the circRNA and (2) the protein cofactor for the noncoding Che-cirC2410 and zma-miR399e5P. Additionally, it is necessary for constructing a genetically stable maize plant to study the pathways participated by zma-mir399e-5P. All these findings may serve as tools in crop disease resistance breeding.

## Materials and Methods

### Pathogenic fungi strains and host plants

The *C. heterostrophus* race O strains DY and WF were collected from previously identified *C. heterostrophus* race O library(3). A GFP coding region, GPDH promoter, and TrpC terminator were joined and constructed into a pCAMBIAth1300 plasmid(S6 Fig.) by using an In-Fusion^®^ HD Cloning Kit (Takara Bio Inc., CA 94043, USA). The multiple cloning sites and plasmid constructing strategy are demonstrated in Supplementary Figure 5. An *Agrobacterium tumefaciens*-mediated transformation (ATMT) transformation method(91) was applied to construct the GFP-tagged fungi DY-GFP and WF-GFP. A fluorescence microscope (LEICA DM2500, Leica Microsystems) was used for observation. Two optical filters, namely, (1) eGFP with excitation wavelength of 460-495 nm and emission wavelength of 510-550 nm and (2) red-green with excitation wavelength of 485-560 nm and emission wavelength of 520-610 nm, were used.

The maize inbred lines Mo17 (resistant to *C. heterostrophus* race O) and B73 (susceptible to *C. heterostrophus* race O) were planted in a 30 cm-diameter pot. The green house temperature was adjusted to 30 °C, and the sodium lamp:metal halide lamp ratio was 1:1. The illuminant power was 400 w/m^2^ for 14 h at daytime and 10 h in the dark.

### RNA sampling and cDNA amplification

Conidium stages were collected from the fungi, which were cultured on a PDA medium for 20 days under 20 °C. The conidium concentration was adjusted to 1×10^5^ spores/ml by using Tween 20 (0.5%). The samples were placed on ice to prevent germination. For each strain, 5 μl of the conidium suspension was inoculated into healthy maize leaves at 4-5 leaf stage. At 9 hpi, the scattered mycelium stage was collected by using #1 oil pike carefully. To collect the dense bundles of strong mycelium stage, a scalpel (#15 blade) was used to cut carefully the leaf tissues at the disease-health interface under a stereoscope. The LCM procedure was performed as described by W. H. Tang(11). The cDNA library preparation was performed using Ovation^®^ RNA-Seq System V2 (NuGEN Technologies Inc., CA, USA) in accordance with the instructions. The nucleic acid QC was performed using an Agilent 2100 Bioanalyzer system. The Agilent RNA 6000 Pico Kit was used for the RNA QC, and the Agilent 1000 Kit was used for the cDNA QC (Agilent Technologies). Three biological repetitions were performed for each fungal pathogen stage. The raw data of RNA-seq was uploaded at the Sequence Read Archive (SRA) database (https://trace.ncbi.nlm.nih.gov/Traces/sra/) (access number:PRJNA436064).

### RNA sequencing and data analysis

Paired-end sequencing was accomplished using Solexa RNA sequencing (OE Biotechnology Co., Ltd., Shanghai, China) on an Illumina HiSeq™ 2500 platform. Low-quality data were discarded. The *C. heterostrophus* race O (strain C5) reference genome data was accessed from https://genome.jgi.doe.gov/portal/CocheC5_3/CocheC5_3.download.html (8). Clean reads and reference genome were matched using TopHat2(92). Transcript construction on the basis of the probability model was conducted using Cufflinks(93). Gene expression was calculated using the fragments per kb per million reads method.

### Bioinformatics prediction of noncoding RNA

LncRNA is a type of RNA molecule that is longer than 200 bp and has no protein coding ability. According to the characteristics of lncRNA, the candidate lncRNA was predicted in three steps: (2) clean reads and reference transcripts were compared, and the recorded coding transcripts in the genome.jgi.doe.gov database were discarded; (2) the transcripts that were filtered in the first step and were longer than 200 bps were reserved; and (3) the transcripts with coding potentials were discarded using four software, namely, CPC(94), CNCI(95), Pfam(96), and PLEK(97).

CircRNA was predicted in accordance with the method reported by Memczak(98). Clean reads and reference genome were compared using Bowtie 2(99).

Interactions between circRNA and maize miRNA and mRNA were predicted using the psRNATarget software (http://plantgrn.noble.org/psRNATarget/)(100). The default prediction parameters were as follows: maximum expectation = 3.0, length for complementarity scoring = 20 bp, and UPE ≤ 25. The miRNA sequence sources of maize were obtained from the miRbase database (http://www.mirbase.org)(101). The circRNA-miRNA-target mRNA interaction network was drawn using Cytoscape 3.6.0(102).Weighted gene coexpression network analysis (WGCNA) of the whole transcriptome was preformed according to Zhang(103).

### Nucleic acid treatment of non-RNA-sequencing material

MicroRNA and total RNA were extracted using E.Z.N.A.^®^ Micro RNA Kit (Omega Bio-Tek, Norcross, GA) in accordance with the manual. Reverse transcription of the total RNA and qRT-PCR were performed using the Transcriptor First-Strand cDNA Synthesis Kit and FastStart Universal SYBR Green Master (ROX) (Roche, USA), respectively, in accordance with the manual. The reverse transcription of microRNA was performed using Mir-X™ miRNA First-Strand Synthesis Kit and SYBR^®^ qRT-PCR (TAKARA, Japan). LightCycler^®^ 96 (Roche, USA) was used for the qRT-PCR and data analysis.

The reference microRNA used for microRNA quantitative analysis was *Z. mays* 5S ribosomal RNA (NCBI accession no.: 4055935), the reference gene for the maize mRNA quantitative analysis was *Z. mays* ACT1 (NCBI accession no.: 100282267). The reference gene for the *C. heterostrophus* gene quantitative analysis was ACT1 (NCBI accession no.: AY748990.1). The primer pairs used for the quantitative analysis are listed as follows:

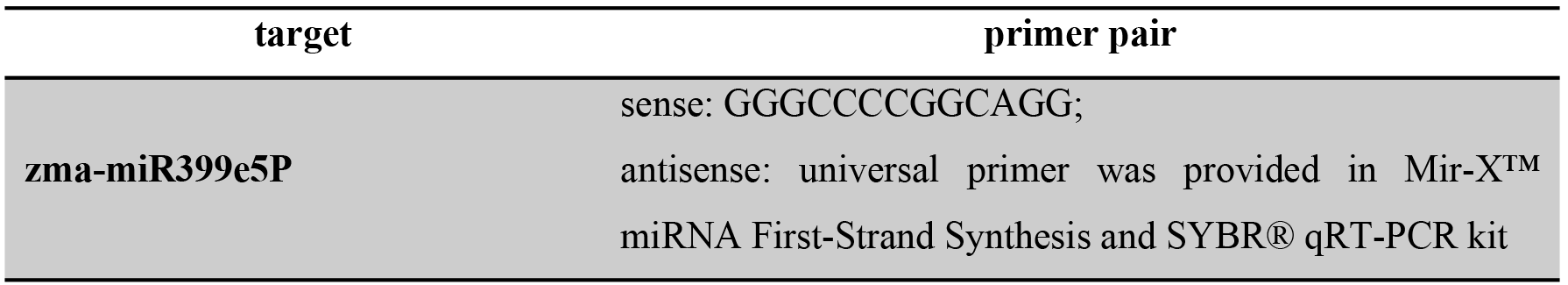

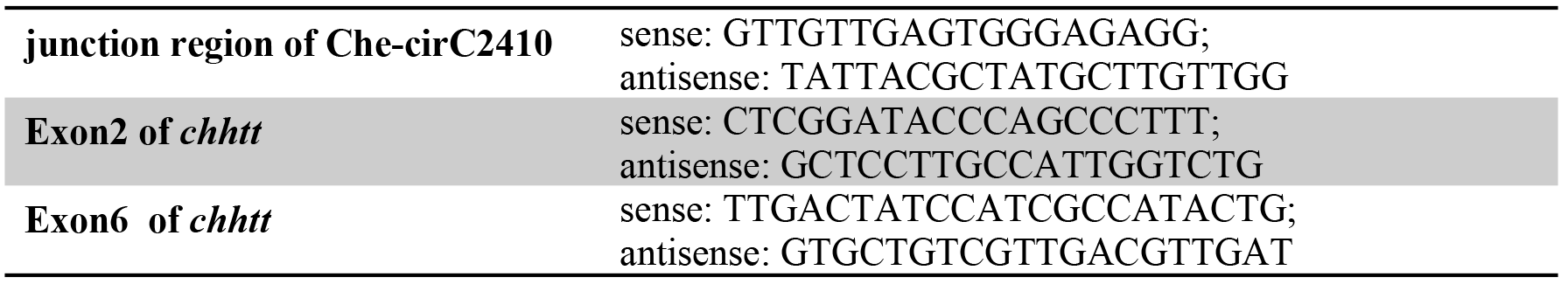

The circRNA treatment by RNase R was performed as described by Ashwal(104) with modifications; the total quantity for digestion was 10 μg, and the digestion time was 1 h.

### In situ hybridization of noncoding RNA

The tissue fixation and paraffin embedding for the in situ hybridization were the same as those of the LCM procedure, except the paraffin was replaced by Paraplast Plus^®^ (0.8% DMSO was added for ease of infiltration and sectioning; Sigma-Aldrich, Merck, Germany). To reduce the RNA loss, the final paraffin penetration cycle was reduced to three rounds. The in situ hybridization probe (BA-F-Che-cirC2410_Junc-C1, 1ZZ) for Che-c2410 was designed to detect the circRNA junction sequence of agtgttgggcgggagaggaggcatggtcagttgtaattaactcttcttcaacctcgagtctagggaacaaatgggtgagccgtgcaggaatggggcggtgtaatgacatggatgtcttgttcttcttcatttggct. The probe was synthesized by Advanced Cell Diagnostics (Advanced Cell Diagnostics Inc., CA 94545, USA); the positive control probe was Probe-BA-Zm-pep1-1zz-st (cat no.: 705101), and the negative control probe was Negative Control Probe-DapB (cat no.: 310043). The miRCURY LNA™ detection probe (product no.: 618532-360) for the zma-miR399e5P in situ hybridization was acquired from Exiqon A/S (Exiqon, Denmark). The positive control probe, /5DigN/AGGATGTTGAGGAAGCGGTCGA/3Dig_N/, was designed simultaneously. The Scramble-miR miRCURY LNA detection probe (product no.: 339111) was used as the negative control. The in situ hybridization of circRNA was performed using the BaseScope™ Detection Reagent Kit (Advanced Cell Diagnostics Inc., CA 94545, USA). The in situ hybridization of microRNA was performed using the miRCURY LNA™ microRNA ISH Optimization Kit (FFPE) (Exiqon, Denmark). The coverslip was deprecated during the incubation of the microRNA probe. Hence, the slides were incubated in a Tissue-Tek^®^ Slide Rack from a HybEZ™ Humidifying System to improve the performance and prevent section loss.

### VIGS assay of the maize genes and knocking out the gene of pathogenic fungi

The VIGS knockdown of the maize target genes was performed in accordance with Rong Wang’s method(105). The primer pairs for building the silencing target into plasmid-2bN81 are listed as follows:

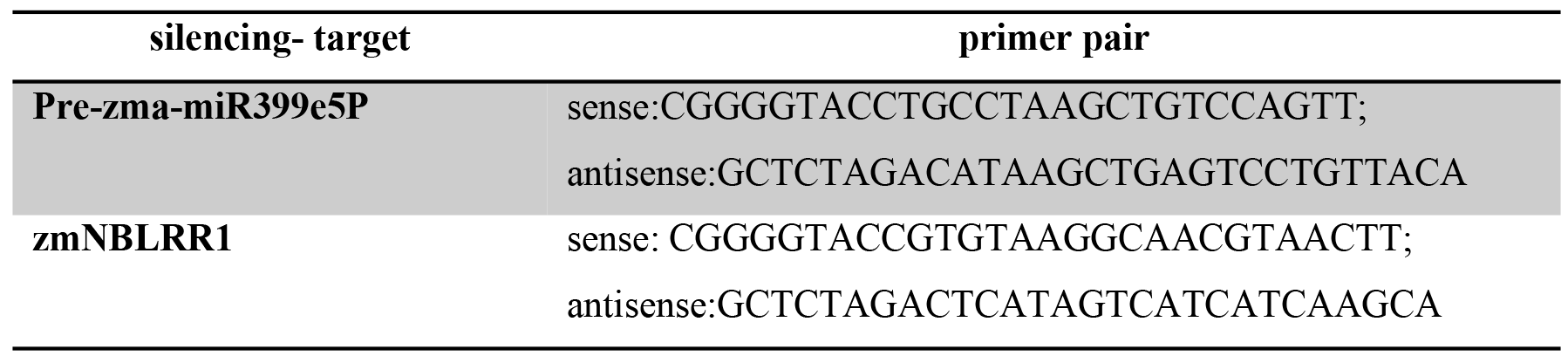

The zma-miR399e5P mimics sequence is GGGCUUCUCUUUCUUGGCAGG; and the antagomir sequence is CCUGCCAAGAAAGAGAAGCCC. All mimics and antagomir were chemically modified by methoxy group, which were finished by Shanghai Sangon Biotech. The mimics and antagomir water solution (Rnase free, 1 nmol/mL) were injected by needleless injector at the separate sides of vein, after this, the pathogen were inoculated as previously described.

ATMT was performed to knock out the *chhtt* gene, which transcribes Che-cirC2410 f in accordance with the method of Gao^45^. The plasmid construction was performed using the In-Fusion^®^ HD Cloning Kit (Takara Bio Inc., CA 94043 USA) and is shown in S6 Fig. The primers for the *chhtt* gene flank regions are listed as follows:

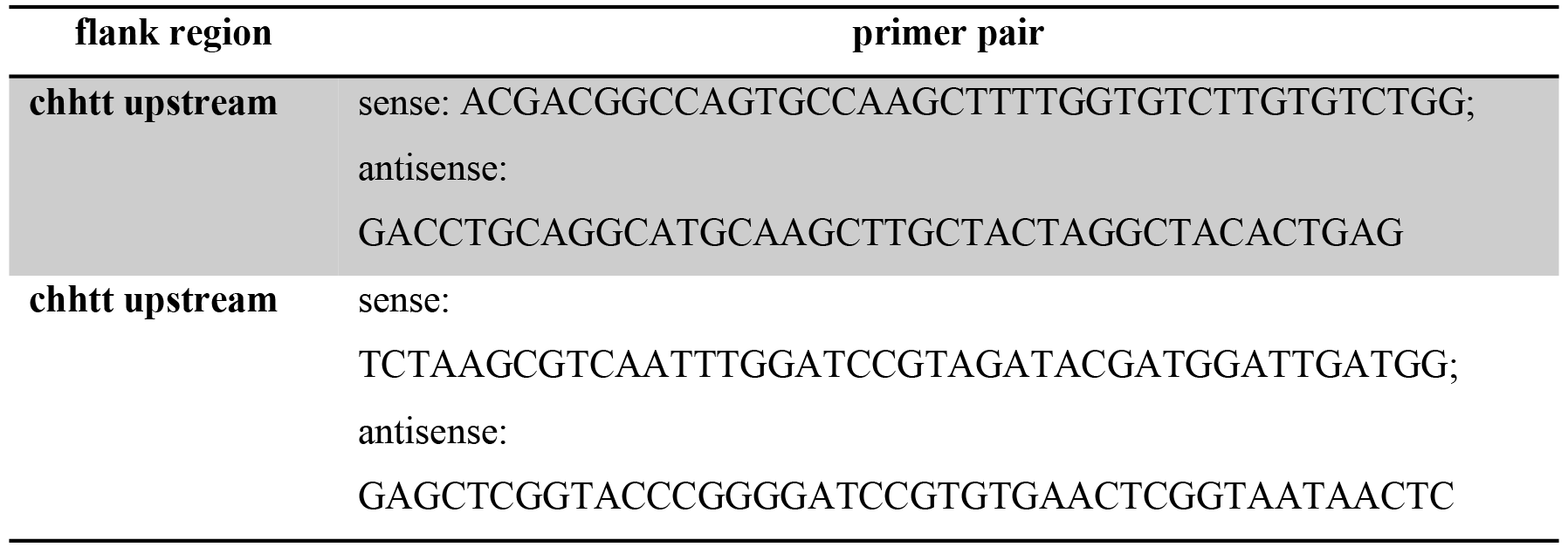

### Dual-luciferase report system

The dual-luciferase report system used was in reference to Runmao Lin’s method(106).

### Infection and disease severity assessment

The infection of *C. heterostrophus* and integral optical density (IOD) calculation were performed as described(3). SPSS 19.0 was used to analyze the significant variations among the IOD values of differential infection. The infection was performed at least thrice.

### Monoclonal antibody, protein treatment, and Western blot

The monoclonal antibody was designed and produced by Youlong Biotech (Shanghai, China). The protein extraction, SDS-PAGE, and Western blot were performed based on the instruction of Burnette(107).

## Acknowledgement

We thank Professor Weihua Tang at Shanghai Institute of Plant Physiology and Ecology for supporting the laser capture microdissection platform, and Professor Tao Zhou in China Agriculture University for supporting the VIGS system in this study.

---

Figure S1.Collection of dense bundles of thick mycelium by using laser capture microdissection (LCM). The left column is the infection complex before LCM collection, and the right column is the infection complex after LCM collection. The top line is the infection complex photographed under a GFP fluorescence channel, and the bottom line is the infection complex photographed under a bright-field channel. MC, motor cell; DM, dense bundles of thick mycelium. Scale bar =75 μm.

Figure S2.Total RNA and cDNA quality control (QC).

(A) QC of total RNA, which is demonstrated by an electrophoretogram on the Agilent 2100 Bioanalyzer system.

(B) QC of cDNA, which is demonstrated by the electrophoretogram.

(C) Chromatographic peak diagram of the total RNA. The sample ID and RNA integrity were labeled on the margin of each diagram in blue color.

(D) Chromatographic peak diagram of cDNA. The sample ID was labeled on the margin of each diagram in blue color

Figure S3.Subcellular location of Che-c2410 and zma-miR399e5P in single species control and mutant. Che-cirC2410 can be detected only in pure *C. heterostrophus*, and zma-miR399e5P can be detected only in pure maize. After the *chhtt* gene was knocked out, Che-cirC2410 cannot be observed anymore. After zma-miR399e5P was knocked down by using VIGS, only a few unsharp dots were detected.

Figure S4.Positive and negative controls to evaluate the BaseScope™ and miRCURY LNA™ ISH. The PEPC can be detected in both the BaseScope™ and miRCURY LNA™ ISH systems; bacterium-dapb (negative control of the BaseScope™ ISH) and scramble-miR (negative control of the miRCURY LNA™ ISH) cannot be detected. The BaseScope™ and miRCURY LNA™ ISH systems are both qualified to analyze the interaction of fungal pathogen and host plants.

Figure S5.The verification of VIGS zma-miR399e5P. The healthy inbred B73 and Mo17 were taken as mock. Four plants of each inbred were introduced for VIGS assay and survey the expression of zma-miR399e5P. After expression test, the lowest expression plant was used for subsequent experiments.

Figure S6.Plasmid construction for GFP tagging and ATMT. .

(A) The pCAMBIA 1300th plasmid was joined with a GAPDH promoter, GFP coding region, and TrpC terminator at the HindIIII and BamHI cloning sites. Kanamycin resistance was used to screen a positive cloning bacterium (Escherichia coli, strain DH5α), and hygromycin B was used to screen a positive fungal mutant.

(B) The pCAMBIA 1300qh plasmid was joined with the upstream and downstream regions of chhtt at multiple cloning sites separately. Kanamycin resistance was used to screen a positive cloning bacterium (E. coli, strain DH5α), and hygromycin B was used to screen a positive fungal mutant.

Table S1.The concentration of RNA, integrity of RNA, cDNA yield and RNA-seq data size. The data reflect the quality control of each steps, from RNA extraction(RNA concentration and integrity), libraries building for sequencing (cDNA concentration, cDNA yield) to RNA-seq (raw reads, filtered reads and filtered bases)

Table S2.Genes revealed by co-expression analysis and fitted in PHI database at the same time. The gene ID conform with the JGI Genome Portal (https://genome.jgi.doe.gov/CocheC53/CocheC53.home.html), the co-expression law is represented by STC code, the function can be found in the gene ontology and annotations lists.

Table S3.The record of circRNA and lncRNA in this report. In the circRNA sheet, information about the genome position, aligned transcripts, annotation of transcripts and transcripts function were exhibited. In the lncRNA sheet, only mean fpkm of each stage was supplied, for the reason that all lncRNAs were novel transcripts.

Table S4.Data set of dual-luciferase reporter system.

S1_dataset. Eighteen interaction matrices including pathogen circRNAs, host miRNAs and host targets.

